# Autonomous multimodal agents enable transparent, spatiotemporal reconstruction of immune dynamics in pancreatic cancer progression

**DOI:** 10.64898/2026.04.20.719684

**Authors:** Beibei Huang, Bo Zhu

## Abstract

Pancreatic cancer progression is orchestrated by dynamic shifts in immune and stromal cellular ecosystems, yet the temporal and spatial principles governing these transitions remain poorly understood. Here, we present an agentic computational pathology framework that leverages large language models to orchestrate modular biomarker inference and spatiotemporal reasoning directly from routine H&E histology. Our approach, ROSIE (RObust in Silico Immunofluorescence), combines deep-learning-based multiplex inference with LLM-driven agent logic that emulates pathologist-level reasoning, enabling transparent and reproducible analysis of complex tissue microarchitectures.

Applying this workflow to pancreatic intraepithelial neoplasia (PanIN) progression in KSC transgenic mice (n=24, ages 4–12 weeks), we generated 10.44 million single-cell profiles and identified a temporally ordered immune trajectory comprising three spatially distinct immune-stromal states: (1) early immune-surveillance niche: sharply bounded window of adaptive immune activation and antigen-presentation enrichment; (2) transitional mixed state: declining lymphoid activity, emerging exhaustion programs, and early EMT/angiogenesis signals; (3) stromal-dominant terminal state: fibroblast expansion, vascular remodeling, and immune silence.

These findings establish pancreatic cancer progression as a temporally ordered sequence of immune activation, exhaustion, and stromal takeover. The agentic framework transcends static AI models by offering dynamic, tool-augmented reasoning that bridges high-dimensional tissue data with clinical interpretability—providing a scalable foundation for identifying therapeutic inflection points in early tumor evolution.

## Introduction

Pancreatic cancer remains one of the most lethal malignancies, driven in part by the clinically silent evolution of precursor lesions that undergo extensive immune and stromal reprogramming long before overt malignant transformation^1–4^. Although early inflammatory activation and progressive fibroinflammatory expansion have been described, **the temporal and spatial mechanisms** that coordinate this transition remain incompletely defined and are an active area of investigation^5–10^.

High-dimensional assays such as single-cell sequencing and multiplexed imaging have revealed key aspects of tumor–immune interactions, but their cost, throughput limitations, and tissue requirements make them impractical for cohort-scale or longitudinal studies^11–18^. In contrast, routine H&E slides are abundant, clinically embedded, and capture the full tissue architecture, yet their molecular content remains largely untapped^19,20^.

Recent advances in computational pathology demonstrate that deep learning can extract latent molecular, architectural, and spatial information directly from H&E images^14,21–23^. Yet despite these capabilities, existing approaches lack an integrated framework capable of reconstructing cell-state transitions, pathway remodeling, and niche-level organization across developmental time^24–27^. No current computational pathology method models temporal progression, and although evolutionary and spatial-omics studies underscore the importance of temporally resolved tissue analysis, they do not provide H&E-based solutions^7,12^.

At the same time, most computational pathology pipelines remain single-task systems, optimized for isolated objectives such as weakly supervised WSI classification, mutation prediction, or spatial feature extraction^1,19,25,28^. These models lack the modularity, transparency, and cross-stage coordination required for large-scale, multi-timepoint tissue analysis. As a result, current workflows cannot unify biomarker inference, spatial organization, and temporal dynamics within a single reproducible framework^7,12,29^. Parallel developments in large language models (LLMs) highlight the potential of agent-like orchestration for complex biomedical tasks, enabling dynamic code generation, modular execution, and multi-step reasoning^30–32^, yet such capabilities have not been systematically applied to computational pathology. Parallel work in the LLM ecosystem shows that agentic toolchains and state-driven orchestration frameworks are still actively evolving and have not yet been systematically applied to computational pathology^33,34^. Together, these gaps motivate the need for a flexible, node-based architecture capable of coordinating heterogeneous analytical modules while maintaining interpretability and scalability across multi-timepoint datasets.

Here, we introduce an ***LLM-orchestrated*** *computational pathology framework* that integrates multi-timepoint H&E slides into high-resolution temporal and spatial maps of pancreatic cancer progression. Using ROSIE-based biomarker inference and modular LLM-driven nodes for feature extraction, time-stratified analysis, and spatial niche mapping, we analyzed 10.44 million cells from KSC mice spanning 4–12 weeks. This framework reveals a coordinated immunological trajectory—from early immune-active niches to mixed immune–stromal states and ultimately to stromal-dominant, suppressive microenvironments—providing a scalable and clinically translatable approach for dissecting early tumor evolution and identifying actionable windows for interception.

### Research Highlights

- **Reconstruction of a stage-resolved immune trajectory in pancreatic cancer.** Using 10.44 million H&E-derived single-cell profiles, we delineate a previously uncharacterized temporal sequence in which early precursor lesions exhibit a sharply defined window of adaptive immune engagement that collapses as disease progresses.
- **Discovery of spatially organized early immune-surveillance niches.** Early lesions contain compact, lymphoid-rich niches with strong antigen-presentation and activation signatures, revealing that early immune recognition is anatomically focused rather than diffuse.
- **Identification of a transitional immune–stromal mixed state.** Intermediate lesions display mixed immune–stromal niches marked by declining lymphoid activation, rising exhaustion programs, and emerging fibroblast- and epithelial-driven EMT/angiogenesis activity—representing a critical inflection point in immune collapse.
- **Characterization of a stromal-dominant, immune-silent terminal state.** Late-stage lesions are dominated by fibroblast-rich niches with high EMT, angiogenesis, fibrosis, and proliferative activity, reflecting a fully reprogrammed microenvironment permissive for invasive carcinoma.
- **H&E-based reconstruction of temporal and spatial microenvironmental evolution.** By integrating ROSIE-based biomarker inference with an **LLM-orchestrated, multi-node computational pathology framework**, we achieve the first H&E-only reconstruction of pancreatic cancer’s temporal immune remodeling and niche-level architectural transitions at scale.

## Results

### 2.1 An LLM-orchestrated framework enables modular spatial and temporal tissue analysis

To enable scalable, high-resolution interrogation of pancreatic cancer progression, we developed an LLM-orchestrated computational framework that integrates multiple computational nodes into a unified analysis pipeline. The dashboard provides an interactive interface for loading whole-slide images (WSIs), executing node-specific tasks, and visualizing intermediate outputs. Each node performs a discrete function—WSI tiling, feature extraction, cell detection, cell typing, temporal modeling, and spatial niche analysis—allowing transparent inspection of data flow and facilitating reproducible, modular experimentation. Representative patches displayed within the dashboard illustrate how raw images, feature maps, and cell-level annotations are surfaced at each stage of the framework.

### 2.2 Multi-node LLM-agent pipeline enables scalable feature extraction and cell-level annotation

We next characterized the performance of the multi-node pipeline underlying the LLM-orchestrated framework. Node 1 partitions WSIs into standardized 512×512 patches using adaptive grid tiling and tissue detection. Node 2.1 applies ROSIE to infer 50-channel protein feature maps and compute spatial coordinates for each detected cell. Node 2.2 performs HED-based nucleus enhancement, watershed segmentation, and region-property extraction to generate per-cell morphological descriptors. Together, these nodes produced a dataset of approximately 10.44 million cells from KSC mice aged 4–12 weeks, providing a dense and spatially resolved representation of the evolving tumor microenvironment. Node 3 integrates these annotations to construct a Timed Petri Net model of cell-state transitions, while Node 4 performs spatial niche analysis by combining cell identities with DBSCAN clustering and pathway scoring. This modular architecture enables end-to-end analysis from raw histology to temporal and spatial modeling.

### 2.3 Temporal shifts in major immune and stromal populations across 4–12 weeks

We first examined how major cell populations change across disease progression. Because structural compartments dominate the tissue composition and obscure low-abundance immune subsets in a single stacked representation, we visualized structural and immune populations separately in Figure 3. Structural cells—including epithelial cells, fibroblasts, and endothelial cells—accounted for the majority of all detected cells and displayed clear temporal trends (**Figure 3A**). Epithelial cells increased steadily from early to late stages, consistent with progressive tumor expansion, whereas fibroblasts peaked at early–mid stages before gradually declining. Endothelial cells remained a minor but stable component throughout.

Immune populations were plotted on an expanded 0–5% scale to resolve their temporal dynamics (**Figure 3B**). Early lesions (4–6 weeks) exhibited substantial lymphoid infiltration, including CD4⁺ T cells, CD8⁺ T cells, B cells, NK cells, and dendritic cells. Myeloid populations—including CD68⁺ macrophages and monocyte/MDSC-like cells—expanded progressively through mid-stage disease, coinciding with increasing stromal burden. These coordinated shifts indicate a transition from an immune-engaged early microenvironment to a more myeloid-dominated and stroma-rich state as lesions advance. In total, we analyzed approximately 10.44 million cells across all time points, with their distributions summarized in **Figure 3**.

### 2.4 Dynamic change rates reveal stage-specific immune remodeling

To quantify temporal transitions in cellular composition, we computed log2 fold changes (LFC) between consecutive timepoints. The resulting heatmap revealed distinct, stage-specific fluctuations across both immune and structural compartments. Early transitions were marked by a pronounced increase in Neutrophils at 4→5 weeks (LFC = 3.15), accompanied by coordinated rises in multiple myeloid subsets, including CD68⁺ myeloid cells and CD163⁺ macrophages. NK cells showed a transient expansion at 5→6 weeks (LFC = 1.72), followed by a reduction at 6→7 weeks (LFC = –1.56).

Intermediate intervals displayed mixed remodeling patterns. At 7→8 weeks, CD68⁺ myeloid cells exhibited a notable increase (LFC = 1.35), whereas the 8→9-week transition showed broad decreases across lymphoid subsets, including CD8⁺ T cells, CD4⁺ T cells, B cells, and NK cells (LFC range: –0.89 to –1.52). Endothelial cells underwent the largest structural shift during this interval (LFC = –4.68).

Later transitions (9→11 and 11→12 weeks) showed partial rebounds in several lymphoid populations, including CD4⁺ and CD8⁺ T cells, along with modest increases in Tregs. Together, these LFC-based measurements delineate discrete temporal phases characterized by coordinated changes in innate, adaptive, and structural cell populations.

### 2.5 Timed Petri Net models the progression of pancreatic cancer across eight developmental stages

To visualize coordinated temporal shifts across cell populations, we constructed a Timed Petri Net using cell-type proportions from 4–12 weeks. Each timepoint is represented as a two-layer node summarizing the dominant structural or immune composition, while transitions encode the magnitude and direction of log2 fold change (LFC) between consecutive stages.

Early intervals were characterized by strong innate immune fluctuations, including a marked increase in Neutrophils at 4→5 weeks and a transient rise in NK cells at 5→6 weeks. Mid-stage transitions showed increases in CD68⁺ myeloid cells at 7→8 weeks, accompanied by broad reductions in multiple lymphoid subsets at 8→9 weeks. Structural populations also shifted during this interval, with Endothelial cells exhibiting the largest decrease across all transitions.

Later stages (9→11 and 11→12 weeks) displayed partial rebounds in several lymphoid populations and modest increases in Tregs. Across all stages, the Petri Net highlights discrete temporal phases defined by coordinated changes in innate, adaptive, and structural compartments.

### 2.6 Global pathway activity dynamics reveal early immune activation and late EMT/angiogenesis programs

To quantify pathway-level remodeling across disease progression, we computed weighted pathway activity scores for eight biological programs using bootstrap-stabilized single-cell profiles from 4–12 weeks. The resulting heatmap revealed a clear temporal architecture. Immune-associated pathways—including Immune Cell Markers, Myeloid Antigen Presentation, and T Cell Activation/Exhaustion—showed their highest activity at 4 weeks (Z = 0.62, 0.35, and 0.33, respectively), consistent with an early, immunologically engaged microenvironment. These signals declined steadily through 9–11 weeks (Z = −0.28 to −0.43), indicating progressive attenuation of immune activation as lesions advanced. Weighted pathway activity was computed as a feature-weighted sum of ROSIE-inferred protein channels, where weights correspond to channel-specific variance contributions estimated from bootstrap resampling.

In contrast, pathways linked to structural remodeling and tumor invasion exhibited delayed but pronounced induction at later stages. EMT activity increased at 11–12 weeks (Z = 0.04–0.21), while Angiogenesis and Fibrosis showed strong late-stage activation (Z = 1.81 and 1.77 at 12 weeks), reflecting stromal expansion and vascular remodeling characteristic of invasive transformation. Pathway range analysis further identified Immune Cell Markers as the most dynamically regulated program (range = 1.04), followed by Myeloid Antigen Presentation and T Cell Activation/Exhaustion, underscoring the centrality of early immune remodeling in shaping tumor evolution. Together, these results delineate a coordinated shift from early immune activation to late-stage stromal and invasive programs.

### 2.7 Spatial niche analysis identifies pathway-level transitions in the evolving tumor microenvironment

To resolve how pathway programs are spatially organized across disease progression, we performed a niche-level analysis integrating CellClassifier-derived cell identities with spatial clustering of cell centroids. The resulting niche-by-pathway profiles revealed a clear temporal architecture that complements the global pathway dynamics in **Figure 6**. Early lesions (4–6 weeks) contained multiple immune-enriched niches characterized by high Immune Activation and Myeloid Suppressive activity, consistent with the lymphoid-dominant microenvironment observed in **Figure 3** and the early activation signatures in **Figure 6**. These immune-active niches were spatially compact and localized around nascent epithelial structures, reflecting coordinated early surveillance.

Intermediate stages (7–9 weeks) showed a marked reorganization of niche structure. Immune-enriched niches exhibited declining activation and rising exhaustion signatures, while fibroblast- and epithelial-rich niches began to display elevated EMT and Angiogenesis activity. This niche-level heterogeneity mirrors the mixed immune–stromal state captured by the Petri Net transitions in **Figure 5** and corresponds to the broad lymphoid contraction and myeloid expansion observed in **Figure 4**.

By late stages (11–12 weeks), the niche landscape was dominated by stromal-rich neighborhoods exhibiting strong EMT, Angiogenesis, and Fibrosis activity. Immune-active niches were largely absent, replaced by spatially coherent fibroblast- and epithelial-driven niches that reflected the late-stage structural remodeling and proliferative surge seen in **Figure 6**. Together, these spatial patterns delineate a coordinated progression from early immune engagement to late stromal and invasive programs, providing a spatially resolved framework for understanding microenvironmental transitions during pancreatic cancer evolution..

## Discussion

### 3.1 A coordinated shift in immune-state dynamics during pancreatic cancer progression

Our findings reveal that immune-state transitions during pancreatic cancer development follow a highly coordinated and temporally ordered trajectory. Early lesions exhibit a distinctly immune-engaged microenvironment, characterized by abundant lymphoid populations—including CD4⁺ and CD8⁺ T cells, NK cells, B cells, and dendritic cells—and strong antigen-presentation and activation signatures. This early immune activation aligns with the sentinel inflammatory responses described in the earlier longitudinal study^35^, which identified TSPO⁺ tumor-associated macrophages as key initiators of early lesion-associated inflammation. However, our single-cell and pathway-level analyses extend these observations by demonstrating that early immune engagement is not diffuse but spatially organized into compact, immune-enriched niches that cluster around emerging epithelial structures.

As lesions progress, this coordinated immune architecture gradually deteriorates. Lymphoid signatures decline, exhaustion programs rise, and the microenvironment becomes increasingly dominated by CD68⁺ myeloid cells, monocyte/MDSC-like populations, and fibroblast-rich stromal compartments. These shifts reflect a broad reprogramming of immune function rather than simple changes in cell abundance. The emergence of suppressive and remodeling states mirrors the global pathway transitions we observe, suggesting that immune decline and stromal activation are tightly coupled processes that unfold in parallel.

By late stages, the microenvironment transitions into a stromal-dominant, immune-silent state characterized by strong EMT, angiogenesis, fibrosis, and proliferative activity. Immune niches become sparse and functionally quiescent, indicating a collapse of coordinated immune surveillance. Together, these findings position immune-state reprogramming as a central axis of pancreatic cancer evolution and highlight how immune activation, suppression, and stromal expansion are embedded within evolving spatial microenvironments that shape the trajectory from precursor lesions to invasive disease.

### 3.2 Early immune engagement and transient activation of anti-tumor programs

Early pancreatic lesions occupy a brief but clearly defined window of heightened immune engagement, during which multiple lymphoid and innate compartments mount coordinated responses to emerging epithelial transformation. Biomarker-defined cell compositions (**Figure 3–4**) reveal that CD4⁺ and CD8⁺ T cells, NK cells, B cells, and dendritic cells dominate the microenvironment at 4–6 weeks, forming the principal immune compartments during the earliest stages of disease. Dynamic change-rate analysis further highlights transient surges in CD8⁺ T cells, dendritic cells, and neutrophils, suggesting rapid recruitment and activation of anti-tumor programs as lesions first arise.

Pathway-level signatures reinforce this interpretation: antigen-presentation, interferon-associated signaling, and immune-activation programs are strongly induced during this early interval, consistent with robust immune surveillance. Spatially, these molecular features are mirrored by the organization of immune-enriched niches that cluster tightly around nascent ductal structures (**Figure** 7), indicating that early immune recognition is not only transcriptionally active but anatomically focused.

Together, these observations suggest that early pancreatic lesions experience a transient phase of coordinated immune activation—one in which lymphoid engagement, antigen processing, and spatially localized surveillance converge. This immune-engaged state is short-lived, however, and precedes the destabilization, exhaustion, and stromal expansion that define subsequent stages of progression.

### 3.3 Progressive erosion of immune activation and emergence of suppressive myeloid programs

As lesions advance beyond the early immune-engaged window, the microenvironment undergoes a coordinated erosion of immune-activation programs accompanied by the rise of suppressive myeloid and stromal compartments. Biomarker-defined cell compositions (**Figure 3**) reveal a steady decline in lymphoid populations—including CD4⁺ and CD8⁺ T cells, NK cells, and dendritic cells—beginning at mid-stage disease, while CD68⁺ myeloid cells, monocyte/MDSC-like populations, and fibroblast-rich stromal cells expand substantially. Dynamic change-rate analysis (**Figure 4**) reinforces this trajectory: early surges in cytotoxic and antigen-presenting populations give way to oscillatory fluctuations and eventual contraction, whereas myeloid and stromal compartments exhibit sustained positive change rates as disease progresses.

Pathway-level remodeling provides further evidence of this transition. Immune-activation and antigen-presentation signatures decline sharply across 7–11 weeks, while fibrosis, angiogenesis, and epithelial–mesenchymal transition pathways rise in parallel (Figure 6). These shifts indicate a progressive dismantling of anti-tumor immune programs and the establishment of a microenvironment increasingly dominated by suppressive myeloid activity and stromal remodeling. Temporal modeling with the Petri Net (**Figure 5**) captures this transition as a shift from lymphoid-driven to myeloid-driven regulatory states, reflecting the loss of coordinated immune surveillance.

Spatial niche analysis (**Figure 7**) situates these molecular and cellular changes within evolving tissue architecture. Mid-stage lesions exhibit mixed immune–stromal niches in which immune compartments lose activation while fibroblast- and epithelial-rich niches begin to acquire EMT and angiogenic activity. By late stages, stromal-dominant niches become the prevailing organizational units, and immune niches become sparse and functionally silent. Together, these findings reveal that immune suppression in pancreatic cancer is not abrupt but emerges through a staged erosion of activation signals coupled with the expansion of myeloid and fibroblast-rich programs that progressively reshape the microenvironment.

### 3.4 Spatial microenvironments shape immune-state transitions

The spatial organization of the tissue microenvironment provides an additional layer of structure to the immune-state transitions observed across disease progression. Niche-level analysis (**Figure 7**) shows that early lesions are dominated by immune-enriched microenvironments characterized by high antigen-presentation activity, strong immune-activation signatures, and close spatial proximity between lymphoid populations and emerging ductal structures. These compact, immune-active niches likely reinforce the early surveillance window described above, enabling coordinated recognition of transformed epithelial cells.

As lesions progress, however, the spatial landscape becomes increasingly heterogeneous. Mixed immune–stromal niches emerge during intermediate stages, reflecting the destabilization of lymphoid programs and the rise of myeloid and fibroblast populations. By late stages, the tissue is dominated by stromal-rich niches exhibiting elevated EMT, angiogenesis, and fibrosis activity—pathway signatures that align with the suppressive myeloid and fibroblast expansion observed in **Figure 3–4** and the pathway remodeling in **Figure 6**. These spatial patterns delineate a clear progression from immune-active → mixed → suppressive/stromal microenvironments, providing a structural basis for the gradual loss of immune activation and the establishment of a permissive niche for tumor progression.

Together, these findings highlight that immune-state transitions in pancreatic cancer are fundamentally spatial processes: early immune-active niches support coordinated anti-tumor responses, whereas late stromal-dominated niches create physical and molecular contexts that favor immune evasion and disease advancement.

### 3.5 Biological and clinical implications for interception and therapeutic timing

The immune-state transitions uncovered by our framework have direct implications for understanding pancreatic cancer evolution and for identifying windows of clinical vulnerability. The early presence of biomarker-defined lymphoid engagement and antigen-presentation activity suggests that precursor lesions are initially embedded within an immune-active microenvironment capable of recognizing and responding to epithelial transformation. This observation is consistent with previously reported biomarker profiles in early pancreatic lesions, where TSPO⁺ tumor-associated macrophages were identified as key initiators of sentinel inflammation^35^. Our results extend this concept by showing that early immune activation is not only detectable but quantifiable at single-cell and pathway resolution, and that it is spatially organized within immune-active niches that closely track emerging ductal abnormalities.

As lesions progress, the coordinated shift toward mixed and ultimately suppressive/stromal microenvironments provides a mechanistic basis for the loss of immune surveillance and the emergence of a tumor-permissive niche. The expansion of CD68⁺ myeloid cells, monocyte/MDSC-like populations, and fibroblast-rich stroma—together with rising EMT, angiogenesis, and fibrosis pathways—suggests that therapeutic strategies targeting myeloid suppression, stromal remodeling, or fibroblast–immune crosstalk may be most effective during this transition phase. Conversely, the early immune-active window identified in **Figure 3–4** and reinforced by niche-level analysis in **Figure 7** may represent an opportunity for immune-modulating interventions aimed at reinforcing endogenous surveillance before suppressive programs become entrenched.

More broadly, the ability of our LLM-orchestrated computational pathology framework to reconstruct high-dimensional immune and stromal landscapes from routine H&E slides carries direct clinical relevance. Because the framework operates on standard FFPE tissue sections without requiring molecular assays, multiplexed staining, or specialized equipment, it is inherently scalable to large retrospective cohorts, prospective surveillance programs, and resource-limited settings. The modular, node-based architecture ensures that individual components—biomarker inference, temporal modeling, spatial niche analysis—can be independently updated or reconfigured as new models or pathological questions arise, establishing a reusable analytical paradigm for spatiotemporal tissue analysis beyond pancreatic cancer. By capturing the spatial and temporal structure of immune remodeling at single-cell resolution, the framework provides a principled basis for identifying high-risk precursor lesions, refining surveillance strategies, and guiding the timing of interception in individual patients. The early immune-active niche identified here—spatially compact, antigen-presentation-enriched, and temporally transient—represents a candidate biomarker signature for lesions that retain immunological vulnerability. Conversely, the shift toward mixed and stromal-dominant niches delineates a prognostic continuum that may stratify patients by proximity to immune collapse and therefore by urgency of intervention. These insights extend the sentinel inflammatory principles established in prior preclinical studies by demonstrating that AI-inferred biomarker landscapes can resolve the earliest stages of pancreatic remodeling with sufficient granularity to support clinical decision-making. Collectively, these results underscore the potential of LLM-orchestrated computational pathology to translate complex spatiotemporal biological signatures into scalable, reproducible, and clinically actionable diagnostic tools—an approach that any laboratory with access to H&E whole-slide images can deploy.

### 3.6 Cross-cancer conservation of the immune trajectory supports a generalized model of microenvironmental collapse

The three-phase immune trajectory identified in pancreatic cancer—early adaptive engagement, transitional immune–stromal mixing, and stromal-dominant immunosuppression—aligns mechanistically with conserved patterns of immune activation and collapse observed across solid tumors. Similar immune-to-stromal transitions, characterized by transient innate immune response followed by myeloid-driven suppression and T cell exhaustion, have been documented in lung adenocarcinoma precursors^36^, suggesting that this progression may represent a conserved hallmark of precancer-to-invasive evolution.

Notably, both systems demonstrate stage-specific vulnerability: early-stage immune-active phases in pancreatic cancer (characterized by CD68+ myeloid expansion and Myeloid Antigen Presentation pathway activity) parallel the TIM-3-enriched innate immune state in lung precancers (AAH stage), whereas late-stage stromal dominance (Fibrosis Z-score=1.77, EMT Z-score=0.21) mirrors the immune checkpoint-driven immunosuppression of invasive lung adenocarcinoma.

This convergence across independent cancer types and assay platforms suggests that immune activation–collapse–stromal takeover may be a generalizable principle of solid tumor evolution, with broad implications for timing therapeutic interception strategies to exploit stage-specific vulnerabilities.

### 3.7 Limitations and future directions

While our framework provides a high-resolution view of immune-state transitions during pancreatic cancer progression, several limitations warrant consideration. First, ROSIE’s biomarker inferences are derived from deep-learning models trained on multiplex immunofluorescence data; although major lineages are robustly predicted, rare or transitional phenotypes may require orthogonal validation using targeted antibody panels or spatial transcriptomics. Second, our analysis is based on cross-sectional sampling of lesions at different developmental stages rather than longitudinal tracking of individual lesions, limiting our ability to resolve clonal evolution or stochastic immune fluctuations over time. Third, although our spatial niche analysis captures the emergence of **immune-active → mixed → suppressive/stromal** microenvironments, the causal interactions among epithelial, myeloid, and fibroblast compartments remain to be experimentally dissected.

Future work integrating longitudinal imaging, perturbation studies, and multimodal spatial assays will help refine the mechanistic basis of immune-state transitions and clarify how specific cellular interactions drive the shift toward suppressive microenvironments. In parallel, expanding ROSIE to incorporate global context features from foundation models may further improve the robustness of biomarker inference across diverse staining conditions. Ultimately, translating these computational insights into clinical workflows will require prospective validation to determine how spatially resolved immune states can guide early detection, risk stratification, and the timing of interception strategies.

## Method

### 4.1 Sample acquisition and tissue preparation

Pancreatic tissues were obtained from **eight H&E-stained whole-slide images (SVS format)** derived from female KSC mice (Ptf1a-Cre; LSL-KrasG12D/+; Smad4fl/fl). These specimens span **4–12 weeks of age**, capturing the continuum from early postnatal precursor lesions to more advanced, stroma-rich disease. This age-structured sampling strategy parallels the disease kinetics established in the corresponding KSC model validation35, in which pancreata exhibit early sentinel inflammation followed by increasing ductal transformation and stromal expansion with age. All slides were generated from **formalin-fixed, paraffin-embedded (FFPE)** pancreatic tissue, sectioned at **5 µm** and stained with hematoxylin and eosin (H&E). After whole-slide preprocessing and quality control, a total of **10,446,317 cells** were segmented and retained for downstream analysis. The eight SVS files constituted the complete dataset for computational analysis; **no KC or wild-type control slides were included**. Each SVS file was processed as an independent whole-slide specimen for downstream biomarker inference, spatial niche analysis, and temporal modeling.

### 4.2 LLM-orchestrated modular computational pathology pipeline

A modular, multi-node computational pathology pipeline was developed, in which Large Language Models (LLMs) orchestrate task execution under human supervision. The pipeline was implemented using **LangGraph** (v1.1.2) ^34^—a state-graph orchestration framework that supersedes linear chain-based approaches by enabling cyclic execution, conditional branching, and persistent state management across nodes—which manages node transitions, data-flow dependencies, and execution state across the multi-node workflow.

This architecture supports an asynchronous development cycle in which nodes can be authored independently, executed chronologically, or dynamically reconfigured to optimize computational flow. Each node is defined through a semantically explicit task template that encodes the user’s intent, ensuring that the operational logic, data dependencies, and expected outputs are unambiguously specified. For instance, whereas the initial pipeline follows a linear path, the structural independence of Node 2.1 (morphological feature extraction) and Node 2.2 (biomarker inference) allows for parallelized execution, significantly reducing total computational latency for large-scale cohorts.

The pipeline comprises the following explicitly defined nodes:

- **Node 1 (Adaptive Tiling):** Performs automated segmentation of SVS files into standardized 512 × 512 patches, employing intensity-based heuristics and tissue-mask refinement to exclude non-informative background regions.
- **Node 2.1 (Spatial Feature Extraction):** Computes per-cell coordinates via a hybrid pipeline of color deconvolution and watershed segmentation, ensuring robust nuclear delineation across diverse tissue contexts.
- **Node 2.2 (Biomarker Inference):** Applies **ROSIE** models to assign cell-type labels and molecular signatures, generating a high-dimensional representation of the tissue microenvironment^23^.
- **Node 3 (Temporal Dynamics):** Aggregates cellular outputs across developmental stages to construct transition models, inferring immune-state trajectories over a 4–12-week continuum.
- **Node 4 (Niche Construction):** Integrates pathway scores with spatial clustering to map molecular programs onto spatially resolved microenvironments.

### 4.3 Literature-Augmented Prompt Generation for Node Execution

To enhance biological specificity in LLM-generated execution scripts, we incorporated domain knowledge from pancreatic cancer literature into the prompt templates at each computational node. We assembled a knowledge base from peer-reviewed publications on pancreatic precursor biology, tumor immune microenvironments, and spatiotemporal tissue remodeling, including the KSC model characterization^35^ and parallel ROSIE-based analyses of lung adenocarcinoma progression^36^.

Relevant passages from this literature corpus were identified and included as biological context in each node’s LLM prompt. This anchored the LLM-generated analysis scripts in established biological priors, improving interpretability and biological coherence across the pipeline.

### 4.4 Human-in-the-loop model selection and execution oversight

To ensure peak inferential accuracy, the workflow incorporates a human-in-the-loop (HITL) model selection layer. During the development phase, researchers can deploy multiple Large Language Model (LLM) variants against the same task template. Each node’s execution logic is generated on-demand by leveraging the LLM’s code-generation capabilities: given a standardized task template, the LLM synthesizes executable Python modules that implement the required computational step. Before execution, these auto-generated modules can be inspected, modified, or validated by the researcher to ensure alignment with analytical intent.

Through the framework’s interactive dashboard, intermediate outputs from these modules are monitored in real time, enabling deterministic comparison of model behavior and empirical selection of the optimal LLM before proceeding to downstream nodes. This modular flexibility—combined with version-locked execution, explicit data-dependency tracking, and human-validated code synthesis—ensures that the transition from exploratory image analysis to high-order niche construction is auditable, scalable, and methodologically rigorous.

LLM model specification and reproducibility. Code generation across all pipeline nodes was performed using two LLM backends evaluated in parallel: **Claude 3.5 Sonnet** (Anthropic, version 20241022) and **Llama 2 (7B)** (Meta AI, quantized GGUF variant served locally via Ollama). Model selection was based on qualitative assessment of generated code across representative data subsets, evaluating syntactic correctness, adherence to task template specifications, and alignment of outputs with expected biological results. Claude 3.5 Sonnet demonstrated consistently higher code quality and was selected as the primary model for all production pipeline runs. Llama 2 (7B) was retained as a lightweight, locally deployable alternative for nodes requiring rapid iteration. All LLM inference was performed with temperature set to 0 to ensure deterministic code output. Task templates used as LLM inputs followed a structured prompt format encoding node-specific objectives, input/output specifications, and execution constraints; representative examples are available from the corresponding author upon reasonable request.

### 4.5 ROSIE biomarker inference

Biomarker-level molecular states were inferred directly from H&E patches using ROSIE, a deep-learning framework trained on paired H&E and multiplex immunofluorescence datasets. For each segmented cell, ROSIE predicted a 50-marker expression profile encompassing immune, epithelial, stromal, and signaling-related proteins23. These inferred profiles were used to classify cells into major lineages (e.g., CD4⁺ T cells, CD8⁺ T cells, NK cells, B cells, dendritic cells, CD68⁺ myeloid cells, monocyte/MDSC-like populations, fibroblasts) and to compute pathway-level scores including antigen presentation, interferon signaling, cytotoxic activity, EMT, angiogenesis, and fibrosis.

All predictions were derived **directly from the eight preprocessed H&E whole-slide images**, without additional staining or molecular assays, allowing scalable, high-dimensional molecular profiling from routine histopathology. This approach provides a unified representation of cellular identity and functional state across the entire developmental continuum.

### 4.6 Validation of ROSIE-Inferred Biomarker Predictions

To validate ROSIE predictions, we compared key computational inferences against orthogonal wet-lab measurements from the same KSC cohort^35^. Across seven biological indicators—immune cell abundance, macrophage polarization, stromal remodeling, and disease staging—ROSIE-inferred profiles were directionally concordant with IHC, trichrome morphometry, and immunofluorescence measurements. Specifically, ROSIE detected progressive CD68⁺ myeloid expansion (log₂ FC = 1.35 at 7→8 weeks) consistent with antibody-based quantification; M2 macrophage dominance and T Cell Exhaustion signatures aligned with CD163⁺/CD80⁺ polarization patterns; and structural inferences (EMT Z = 0.21, Fibrosis Z = 1.77 at 12 weeks) mirrored trichrome-confirmed stromal expansion and acinar loss. These cross-modality validations demonstrate that ROSIE accurately captures cellular and molecular dynamics of pancreatic cancer progression from H&E images alone, supporting the biological validity of computational profiles presented throughout this study. Detailed validation of each biological indicator, including quantitative metrics and comparative figures, is provided in **Supplementary Sections 4.6.1–4.6.5**. A summary of concordance across all seven biological indicators is provided in Supplementary **Table 4.6**”

### 4.7 Temporal modeling (change-rates, Petri Net)

Our workflow does not infer temporal progression from single H&E slides; rather, it organizes multi-timepoint H&E datasets into a coherent temporal structure, enabling quantification of observed biological transitions without performing temporal prediction. These slides represent true chronological timepoints, and temporal analyses in subsequent sections rely on this age-structured sampling rather than temporal inference from single images.

To characterize immune-state dynamics across the 4–12-week developmental window, we modeled temporal changes in both cell populations and pathway activities. For each cell type and pathway, we computed **discrete change-rate matrices** that quantified increases, decreases, or oscillatory behavior between adjacent timepoints. These matrices enabled systematic identification of coordinated transitions, including lymphoid contraction, myeloid expansion, and progressive stromal remodeling.

To capture higher-order regulatory structure, we constructed a **timed Petri Net** representing transitions among immune and stromal states. Nodes corresponded to biomarker-defined cell populations, and directed edges encoded inferred transitions derived from the change-rate patterns. Transition timing was anchored to developmental stage, allowing reconstruction of the shift from early immune-active states to intermediate mixed states and ultimately to suppressive/stromal-dominant states. This Petri Net provided a compact, interpretable model of immune-state evolution across the precursor-to-invasive continuum, integrating both temporal ordering and regulatory directionality.

### 4.8 Spatial niche analysis

Spatial microenvironments were inferred by integrating ROSIE-derived pathway scores with **per-cell spatial coordinates** obtained from the segmentation pipeline. For each whole-slide image, DBSCAN was applied to the set of detected cell centroids to group spatially contiguous cells into preliminary clusters. These clusters were subsequently refined using K-means merging to stabilize niche boundaries and ensure consistent granularity across slides. Each resulting niche was assigned a molecular signature based on aggregated pathway scores and cell-type composition, enabling classification into immune-active, mixed immune–stromal, or suppressive/stromal categories.

Niche-level maps were generated for all eight SVS slides, revealing a consistent spatial trajectory across disease progression. Early lesions were dominated by compact immune-active niches enriched for antigen-presentation and immune-activation programs. Intermediate stages exhibited mixed niches characterized by destabilized lymphoid programs and emerging myeloid and fibroblast activity. Late lesions were enriched for stromal-dominant niches displaying elevated EMT, angiogenesis, and fibrosis signatures. These spatial patterns provided the structural basis for the immune-active → mixed → suppressive/stromal transitions described in the Results and Discussion, demonstrating that immune-state evolution is fundamentally embedded within the architecture of the tissue microenvironment.

## Data and Code Availability

The **ROSIE** deep-learning framework, including model architecture, training methodology, and inference code, is publicly available at: https://gitlab.com/enable-medicine-public/rosie.

The original CODEX multiplex immunofluorescence datasets used to train ROSIE are subject to licensing restrictions and patient-privacy protections; access may be granted by Enable Medicine under appropriate data-use agreements.

All **processed datasets generated in this study**—including cell-phenotype annotations (14 cell types), inferred 50-marker expression profiles, DBSCAN-derived spatial clusters, pathway-enrichment scores, spatial-niche assignments, and all quantitative values underlying Figures 1–7 and Supplementary Figures S7.1–S7.8—are provided as **Supplementary Data** files.

**Figure 1.**
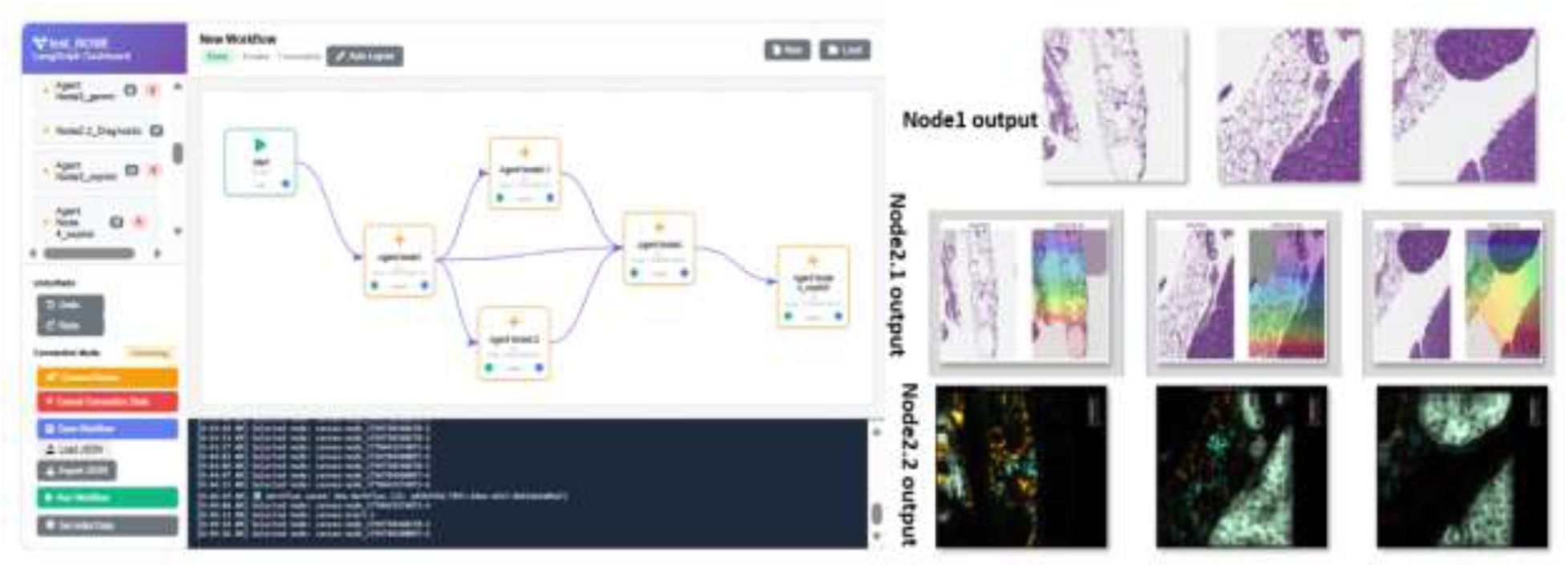
LLM-orchestrated computational pathology framework dashboard for orchestrating the biomarker-exploration pipeline. Shown is the LLM-orchestrated computational pathology framework developed for this study, which provides an interactive interface for coordinating a modular, multi-node workflow for image-based biomarker exploration. The dashboard visualizes node connectivity, execution flow, and intermediate outputs, enabling transparent inspection of each computational stage. On the right, three representative patches illustrate how raw images, feature maps, and cell-level annotations are surfaced during workflow execution. Detailed node-level functions and LLM-generated execution templates are described in Figure 2.

**Figure 2.**
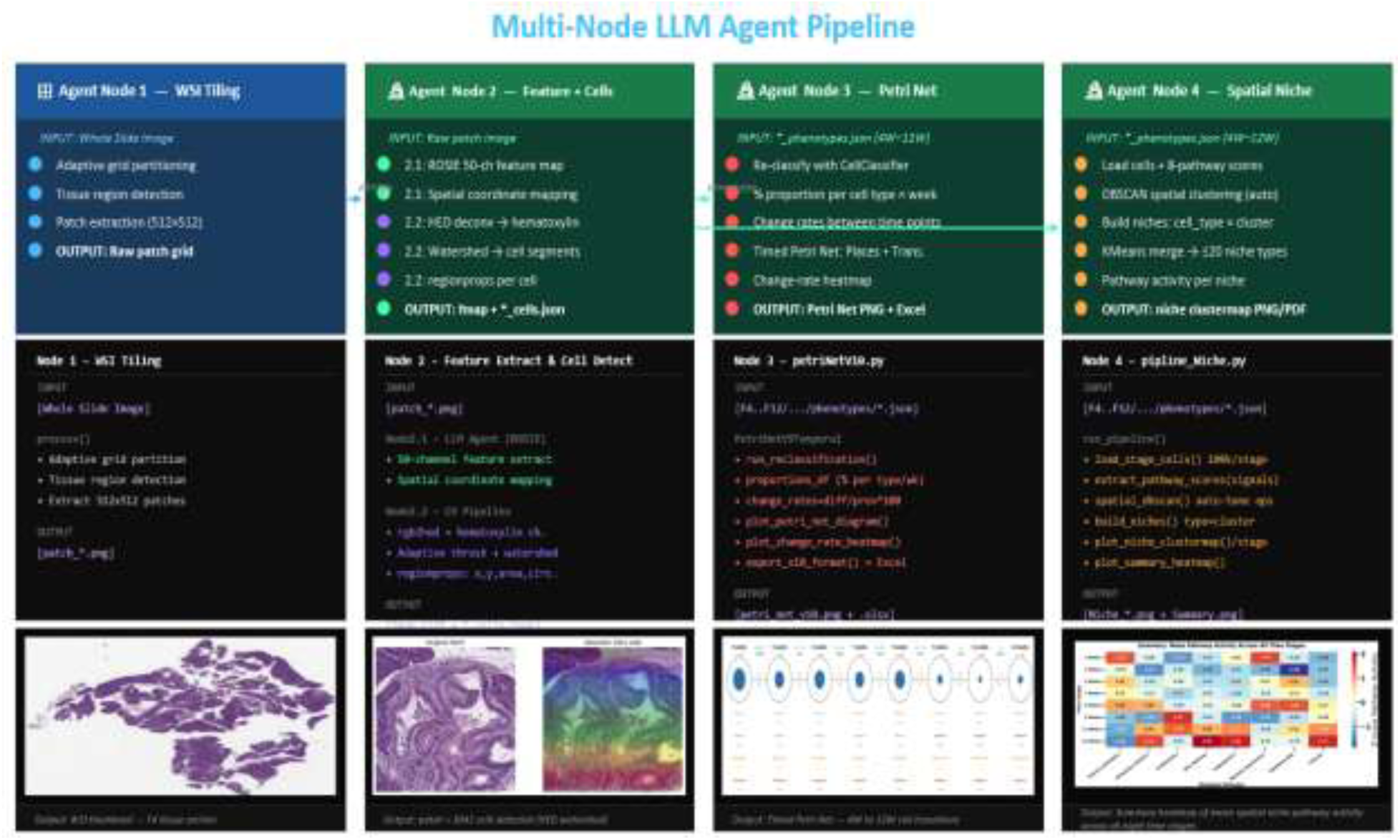
Multi-node LLM-agent pipeline for spatial and temporal tissue analysis. A four-node modular workflow transforms whole-slide images into multiscale biological readouts through LLM-generated execution plans. For each node, a structured template is provided to an LLM agent, which synthesizes a task-specific execution script and triggers downstream computation. **Node 1** receives a WSI and, using an LLM-generated tiling plan, performs adaptive grid partitioning, tissue-region detection, and extraction of standardized 512×512 patches. **Node 2** integrates two LLM-driven submodules: Node 2.1 generates a ROSIE-based feature-extraction script that produces 50-channel protein feature maps and spatial coordinates, while Node 2.2 generates a computer-vision execution plan for HED deconvolution, adaptive thresholding, watershed segmentation, and per-cell morphological quantification. **Node 3** loads phenotype JSON files across developmental stages (4–12 weeks) and uses an LLM-generated temporal-modeling template to reclassify cells, compute cell-type proportions and change rates, and construct a timed Petri Net describing state transitions. Reclassification does not alter biological cell identities; instead, it harmonizes stage-specific annotation noise by applying a probabilistic smoothing model across adjacent timepoints. **Node 4** applies an LLM-generated spatial-analysis plan to integrate pathway scores with DBSCAN clustering and KMeans merging, producing niche-level pathway-activity heatmaps that summarize microenvironmental remodeling. **The third row shows representative examples of the outputs generated by each node’s execution.** Together, these nodes demonstrate how LLM-generated execution plans can orchestrate a fully automated, interpretable pipeline for spatial, temporal, and pathway-level tissue analysis. Representative multi-panel ROSIE biomarker predictions (Node 2.1) and cell-detection visualizations (Node 2.2) are provided in Supplementary Figure S2 to illustrate intermediate outputs of the feature-extraction and segmentation modules.

**Figure 3.**
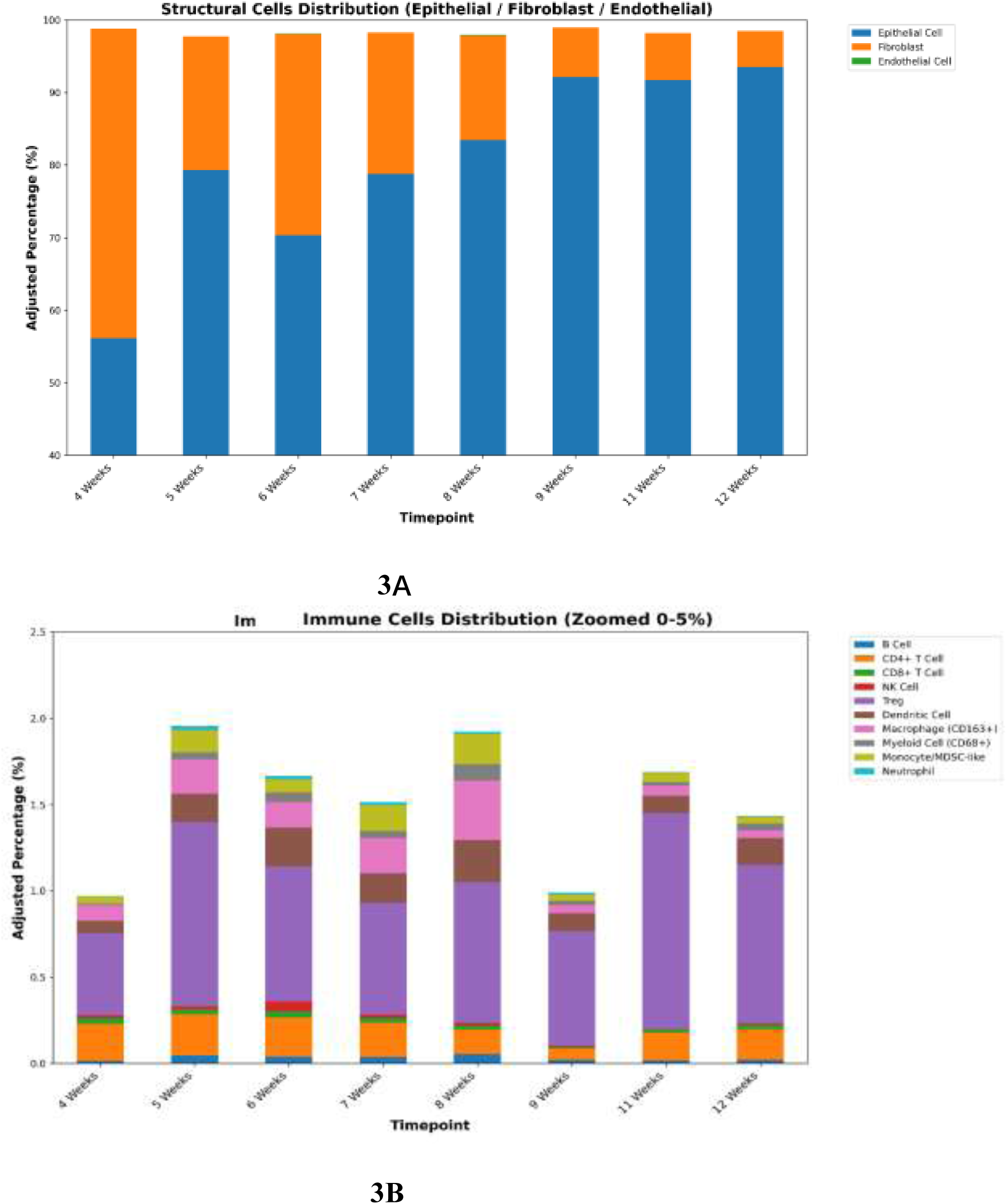
Temporal dynamics of major cell phenotypes during disease progression. (A) Stacked bar chart showing adjusted proportional abundances of structural compartments—including epithelial cells, fibroblasts, and endothelial cells—across eight timepoints (4–12 weeks). These populations dominate overall tissue composition and display clear temporal trends, with epithelial cells increasing steadily and fibroblasts peaking at early–mid stages before declining. (B) Immune populations—including B cells, CD4⁺ T cells, CD8⁺ T cells, Tregs, NK cells, dendritic cells, neutrophils, monocytes/MDSC-like cells, and CD68⁺ myeloid cells—are visualized separately on a zoomed 0–5% scale to resolve low-abundance subsets that would otherwise be obscured by structural compartments. Immune populations exhibit distinct temporal shifts, with early lymphoid enrichment followed by progressive myeloid expansion. Together, these trends illustrate a transition from an immune-engaged early microenvironment to a more myeloid-dominated and stroma-rich state as lesions advance.

**Figure 4.**
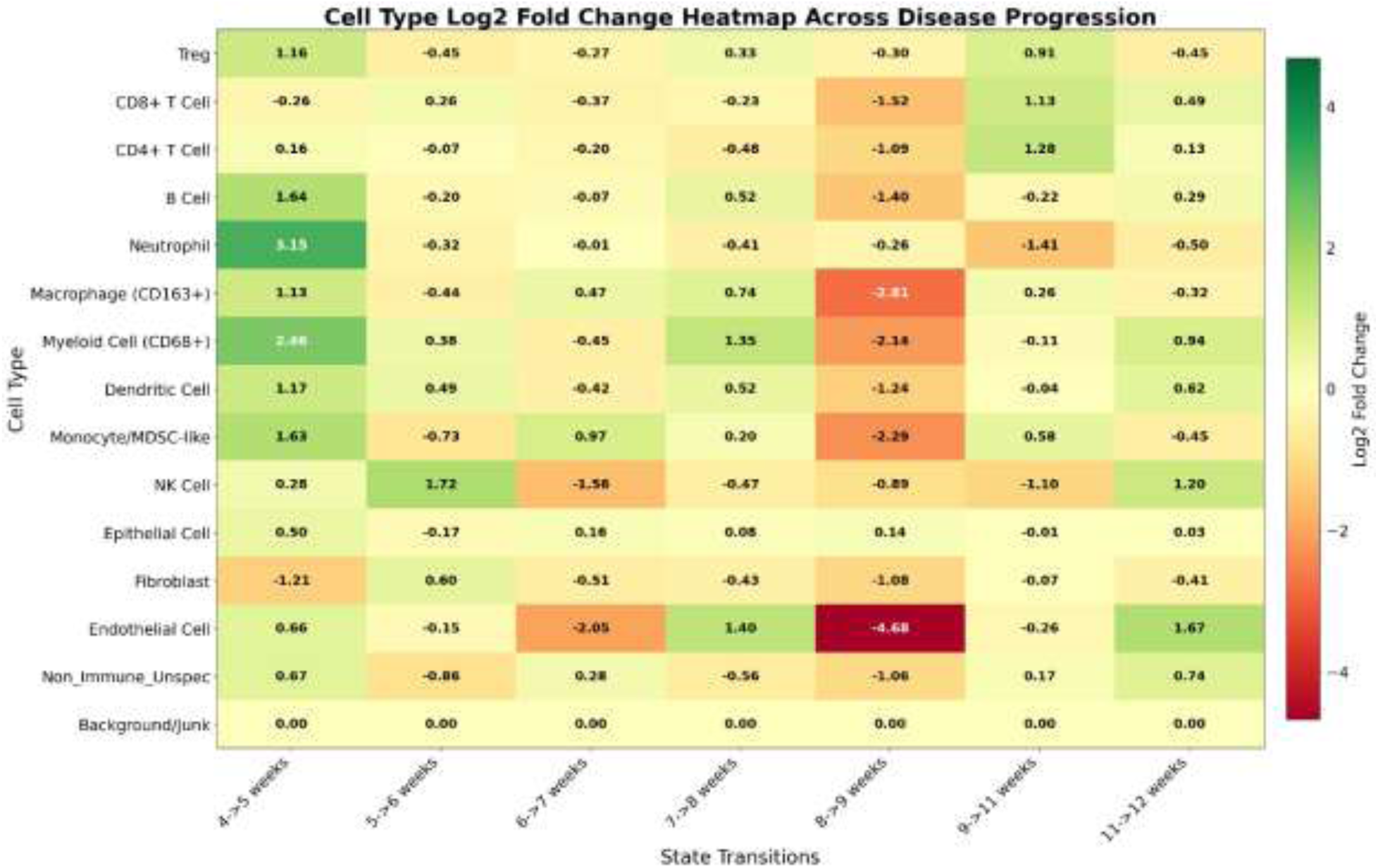
Dynamic change rates of major cell types across consecutive disease-progression intervals. This heatmap summarizes temporal changes in cell-type composition using log2 fold change (LFC) between consecutive timepoints. Each column represents a state transition (e.g., 4→5 weeks), and each row corresponds to a specific cell type. Positive LFC values (green) indicate increased relative abundance, whereas negative values (red) indicate decreases. Major transitions include a sharp rise in Neutrophils at 4→5 weeks (LFC = 3.15), a transient expansion of NK cells at 5→6 weeks (LFC = 1.72), and a marked reduction of Endothelial cells at 8→9 weeks (LFC = – 4.68). These LFC patterns delineate distinct temporal phases of immune and structural population shifts.

**Figure 5.**
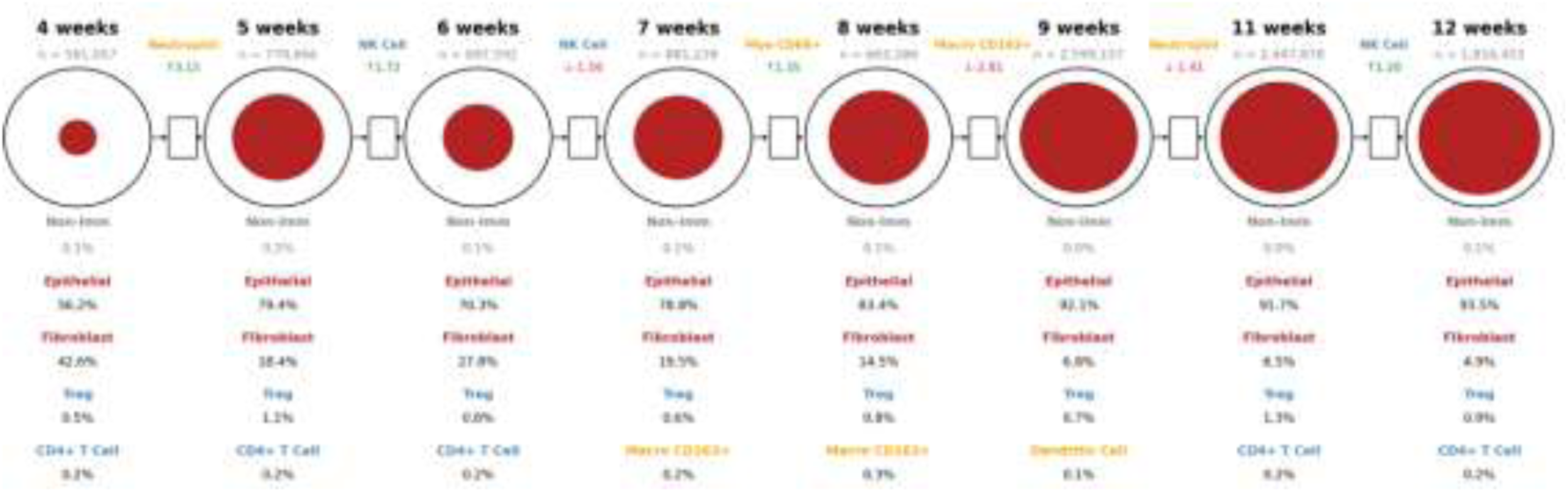
Timed Petri Net summarizes dynamic shifts in major immune and structural cell populations across eight timepoints (4–12 weeks). NK cells are placed at the central node because they function as the dominant upstream regulator of macrophage polarization: Each node represents a timepoint and displays the top four cell types by relative abundance, the non-immune fraction, and the dominant structural or immune lineage. Directed transitions between nodes indicate temporal progression, with annotated log2 fold change (LFC) values marking the cell type exhibiting the largest magnitude change for each interval. Early transitions include a strong increase in Neutrophils at 4→5 weeks and a transient rise in NK cells at 5→6 weeks. Mid-stage transitions show increases in CD68⁺ myeloid cells at 7→8 weeks and broad reductions in lymphoid subsets at 8→9 weeks, including a pronounced decrease in Endothelial cells. Later transitions show partial recovery of several lymphoid populations. Together, the Petri Net provides a stage-resolved view of coordinated shifts across immune and structural compartments

**Figure 6.**
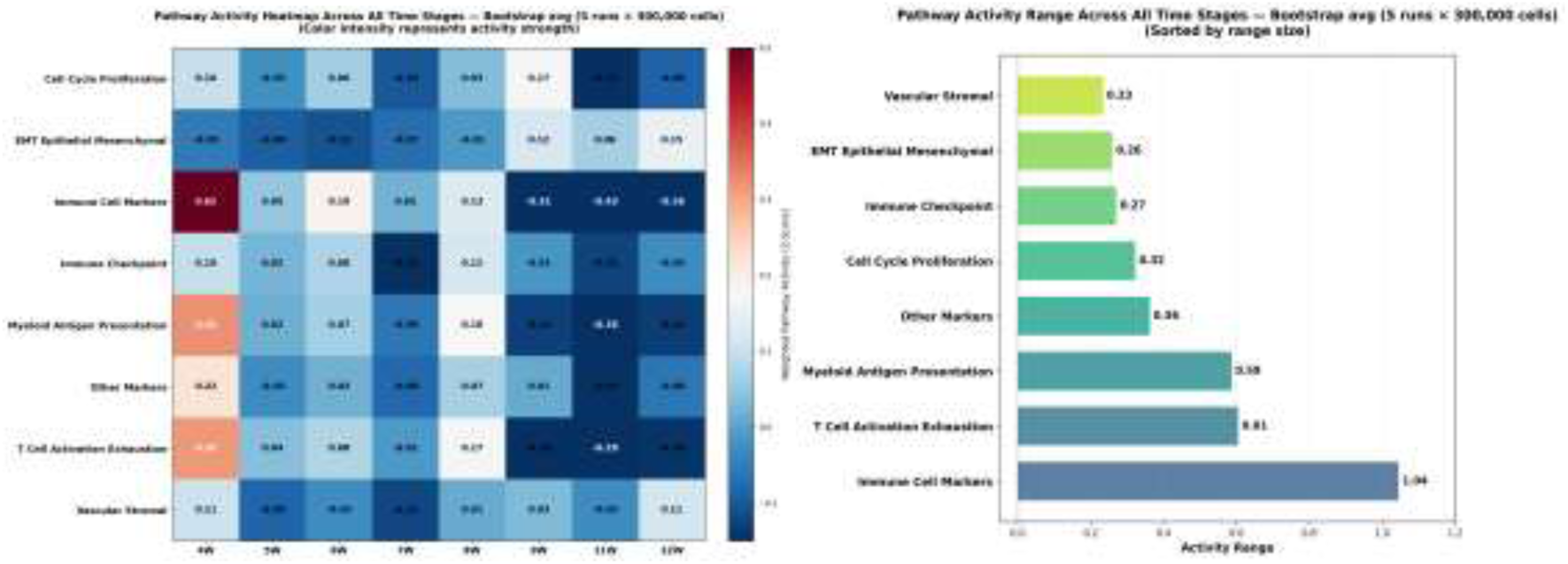
**Left**: Weighted pathway activity (Z-score) heatmap across eight time points (4–12 weeks) in a murine pancreatic cancer progression model, derived from bootstrap-averaged estimates (5 runs × 300,000 randomly sampled cells per stage per run). Immune-associated pathways—including Immune Cell Markers, Myeloid Antigen Presentation, and T Cell Activation/Exhaustion—showed high activity at 4 weeks, followed by a progressive decline through 9–11 weeks, consistent with early immune engagement and subsequent suppression during lesion progression. In contrast, EMT and Vascular/Stromal programs exhibited delayed activation, increasing at 11–12 weeks, reflecting late-stage epithelial– mesenchymal transition and stromal/angiogenic remodeling. **Right**: Corresponding pathway activity ranges ranked by temporal variability. Immune Cell Markers displayed the largest dynamic range (1.04), followed by T Cell Activation/Exhaustion (0.61) and Myeloid Antigen Presentation (0.59), highlighting strong early immune remodeling. Structural pathways such as EMT (0.26) and Vascular Stromal (0.23) showed more modest but late-stage increases. These results collectively delineate a shift from early immune activation to late stromal and invasive programs during tumor evolution.

**Figure 7.**
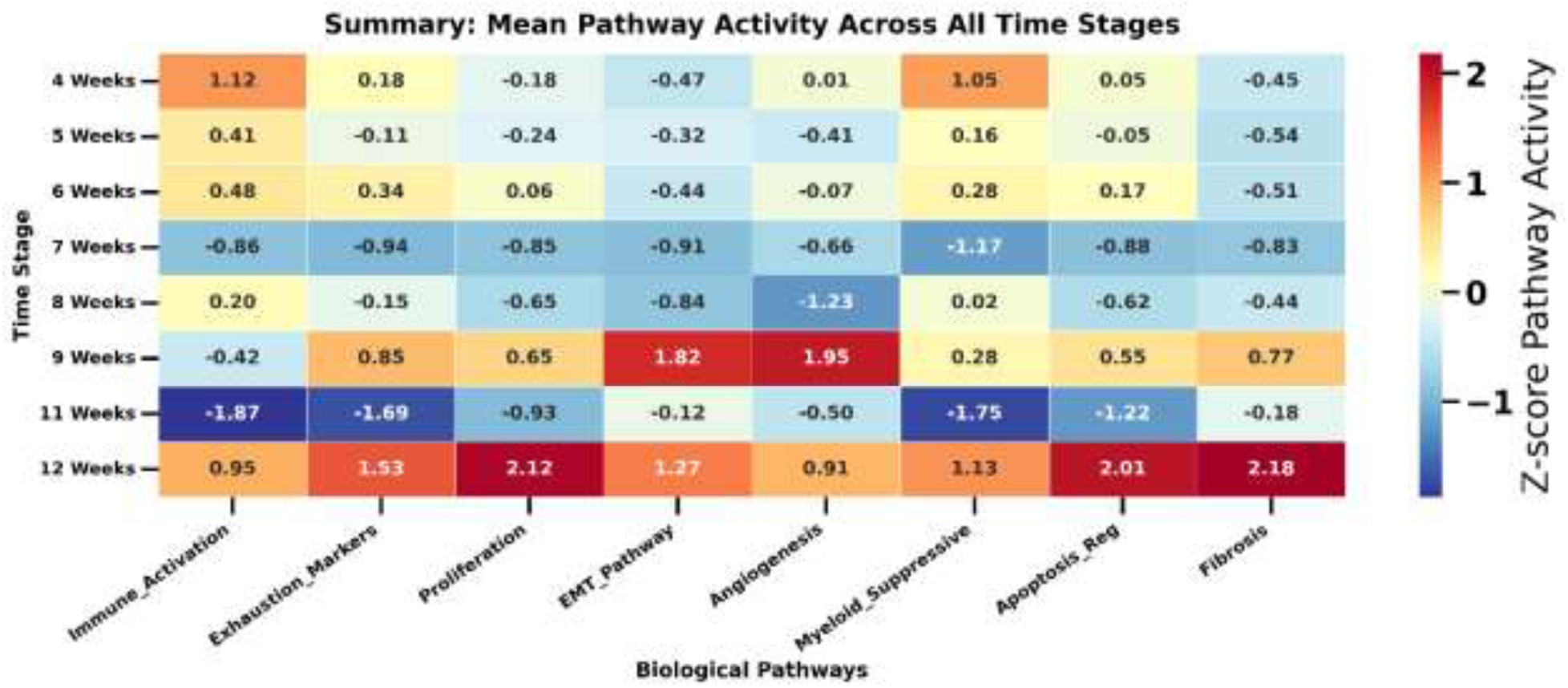
Summary heatmap of mean spatial niche pathway activity across all eight time stages (4–12 weeks) Heatmap showing Z-score–normalized mean pathway activity across all spatial niches identified at each time stage. Each column represents a biological pathway, and each row corresponds to a developmental stage. Early lesions (4–6 weeks) exhibit strong Immune Activation and Myeloid Suppressive activity, reflecting an immune-engaged microenvironment. Intermediate stages (7–9 weeks) show declining immune activity and the emergence of mixed immune–stromal niches with rising EMT and Angiogenesis signals. Late stages (11–12 weeks) are dominated by stromal-rich niches with high EMT, Angiogenesis, Fibrosis, and Proliferation activity, consistent with advanced epithelial–mesenchymal transition, vascular remodeling, and stromal expansion. Stage-specific niche-level heatmaps are provided in Supplementary Figures S7.1–S7.8.

The **raw H&E whole-slide images (SVS format)** used for ROSIE inference were generated from KSC mouse pancreata as described in Section 4.1 and are available from the corresponding author upon reasonable request, subject to institutional guidelines.

The LangGraph-based orchestration pipeline (LangGraphPrj_V5), including the multi-node workflow, node templates, and dashboard interface, is publicly available at: https://github.com/bayjuan5/LangGraphPrj_V5. Custom analysis scripts for spatial clustering, temporal modeling, and pathway analysis are available from the corresponding author upon reasonable request.

## Conclusion

This study shows that routine H&E slides, when processed through a LLM-orchestrated computational pathology framework, can yield a high-resolution, 10-million-cell view of pancreatic cancer evolution. By integrating ROSIE-based biomarker inference with temporal modeling and spatial niche analysis, we uncover a tightly ordered trajectory: early immune-active niches collapse into mixed states and ultimately into stromal-dominant, suppressive microenvironments. These transitions reflect a coordinated loss of adaptive surveillance and the rise of myeloid and fibroblast programs that shape malignant progression. Our results establish a scalable, interpretable framework for extracting molecular and spatial insights directly from standard histology, opening new opportunities for large-cohort studies and for identifying actionable windows for early interception.

## Acknowledgements

We thank the Texas Advanced Computing Center (TACC) for providing computational resources through the project “Predictions of Multi-targeting Drug Response for Colorectal Cancer Using AI Techniques and Single-Cell Analysis” (Grant No. MCB23032). We also acknowledge support from the Texas Workforce Commission. We thank the Manning laboratory for providing the murine tumor samples that were essential for generating the experimental data used in this study.

## Ethics declarations

All animal procedures followed institutional, state, and federal regulations and were approved by the University of Texas MD Anderson Cancer Center Institutional Animal Care and Use Committee (IACUC) prior to implementation. All mice were maintained in filter-topped cages on autoclaved food and water at MD Anderson Cancer Center animal facility and were used in accordance with AALAS and NIH guidelines and regulations.

## Supplementary Information

**Supplementary Figure S2 Intermediate outputs from Node 2.1 and Node 2.2.**

**(A) Representative multiplex ROSIE predictions outputs** from Node 2.1. Multi-panel visualization of inferred 50-channel protein expression across 512×512 pixel patches. Each panel highlights a specific biomarker channel, capturing the spatial heterogeneity and co-localization of predicted proteins across the epithelial, immune, and stromal compartments. Representative patches from two longitudinal timepoints (4 and 12 weeks) are shown to illustrate the progression of microenvironmental remodeling.

**Supplementary Figure S2.1.**
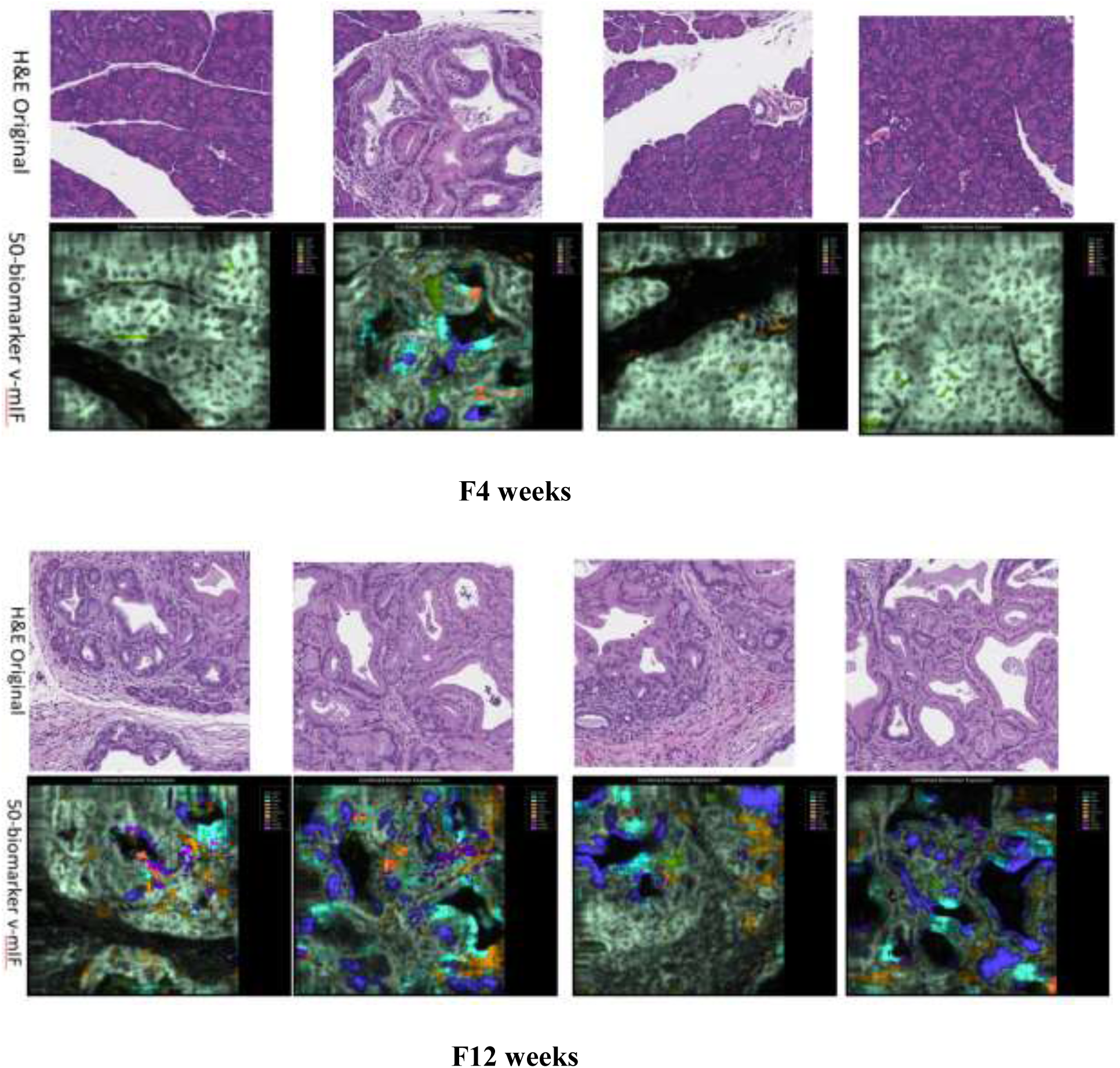
Spatiotemporal mapping of the top 10 inferred biomarker signatures. Multi-panel ROSIE predictions visualize the **top 10 most prominent protein expression profiles** derived from representative 512×512 pixel H&E patches across early (4-week, top) and late-stage (12-week, bottom) lesions. For each timepoint, routine H&E morphology is spatially aligned with its corresponding **virtual multiplex immunofluorescence (mIF)** maps. This high-resolution inference resolves distinct protein gradients and compartment-specific molecular signatures within epithelial, immune, and stromal niches. Collectively, these panels illustrate a systematic microenvironmental shift: transitioning from an immune-active early precursor state to the fibrotic, stromal-dominant, and immunologically cold landscapes characteristic of advanced pancreatic cancer. **(B) Cell-detection and segmentation outputs** from Node 2.2, including HED-enhanced nuclei, watershed-derived cell boundaries, and per-cell morphological annotations. These visualizations demonstrate the accuracy and granularity of the segmentation pipeline used to generate cell-level descriptors for downstream temporal and spatial modeling.

**Supplementary Figure S2.2.**
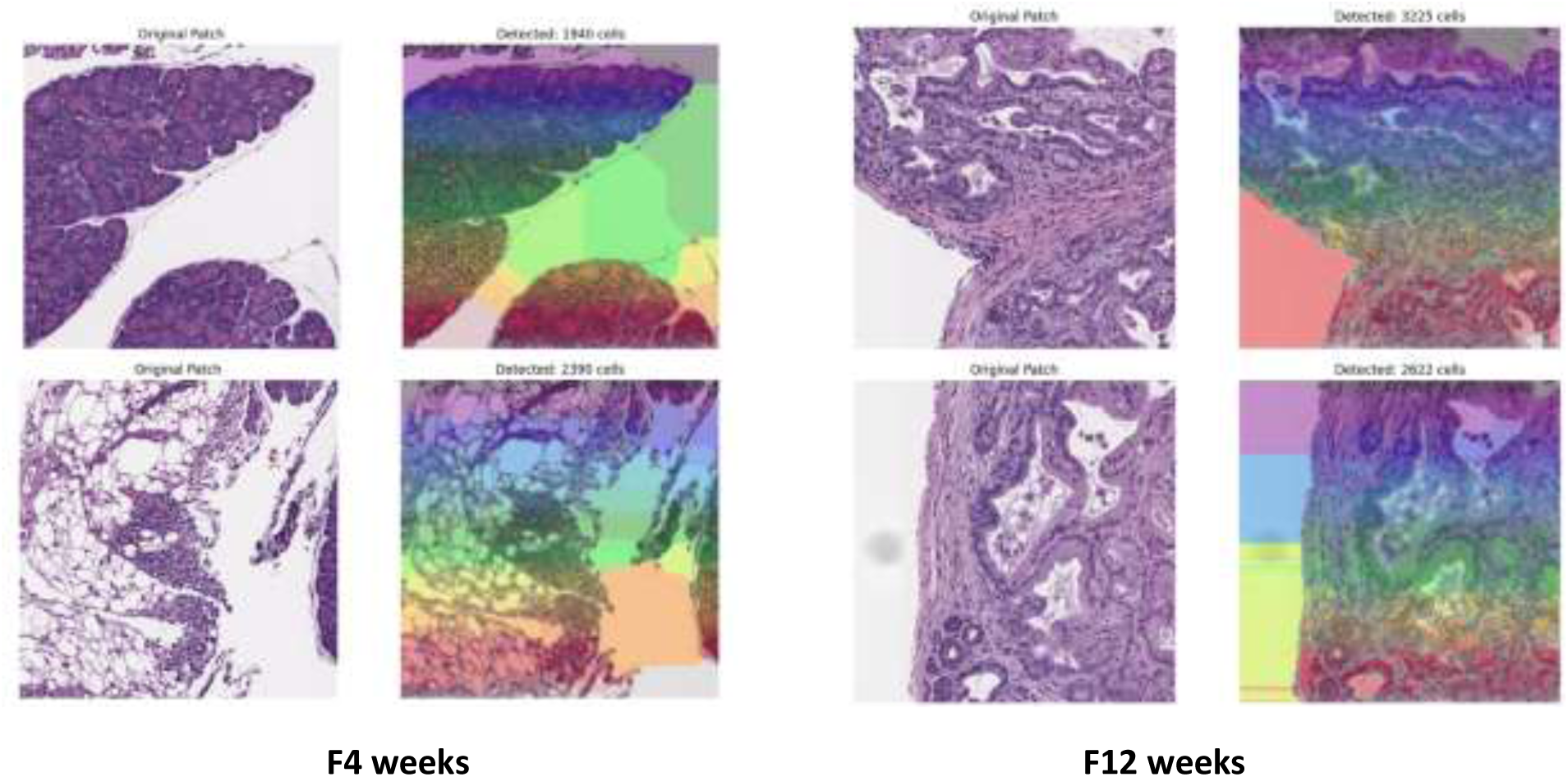
High-fidelity cell segmentation and morphological profiling via Node 2.2. Representative 512×512 pixel H&E patches and corresponding segmentation outputs from early (4-week) and late-stage (12-week) lesions. For each timepoint, original tissue morphology (left) is paired with detected nuclei featuring watershed-derived boundaries and single-cell annotations (right). Node 2.2 integrates HED-based nucleus enhancement with adaptive thresholding to generate precise cell-level masks. This robust segmentation framework provides the foundation for quantifying single-cell density, morphology, and spatial organization, enabling the high-resolution mapping of biomarker expression across tumor progression.

**Figure S7.1.**
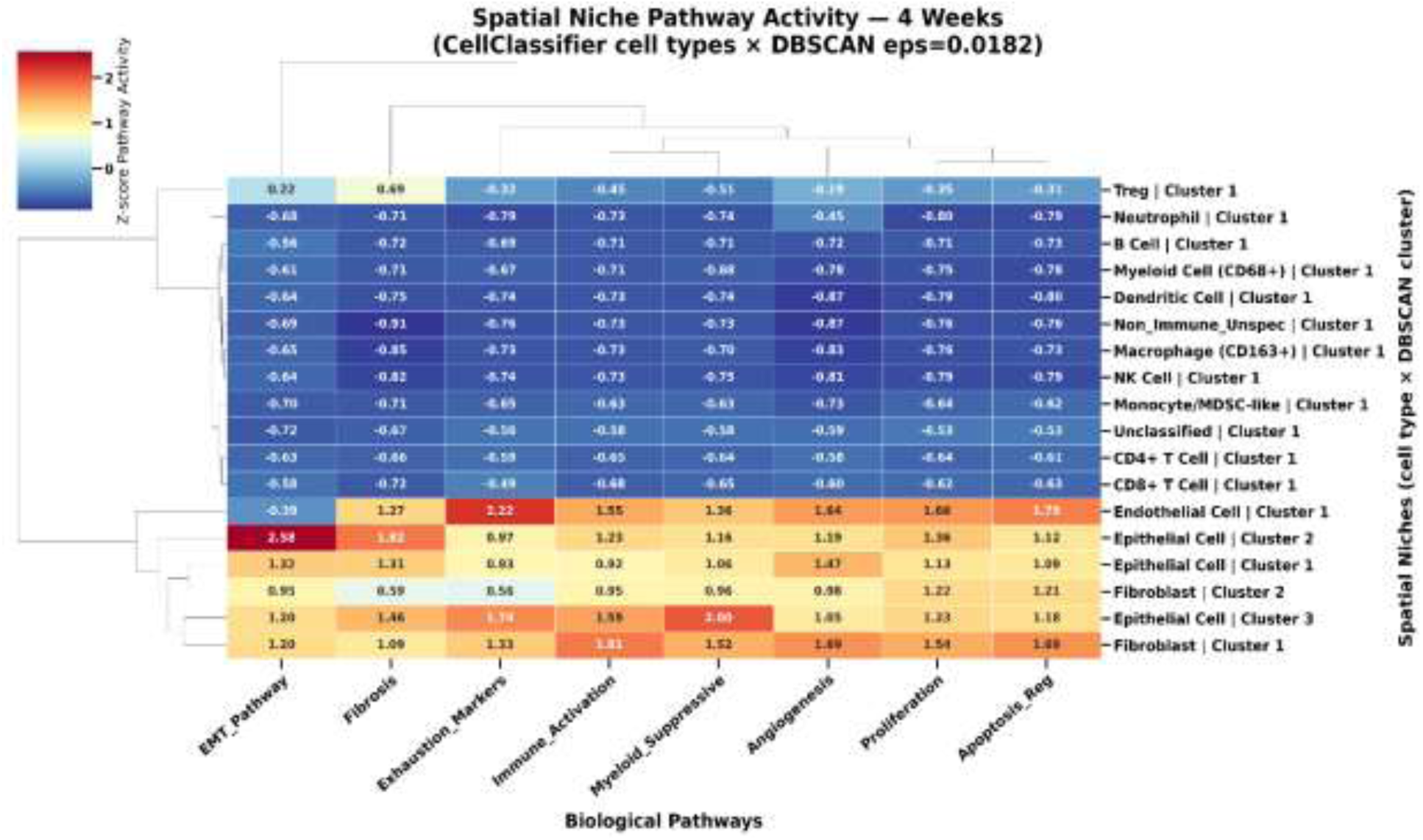
Spatial niche pathway activity at 4 weeks. Heatmap showing Z-score–normalized pathway activity across spatial niches identified at 4 weeks. Each row represents a niche defined by CellClassifier-derived cell identity combined with DBSCAN spatial clustering, and each column corresponds to a biological pathway. Early lesions exhibit multiple immune-enriched niches, including T cells, B cells, dendritic cells, NK cells, and myeloid populations, all displaying elevated Immune Activation and Myeloid Suppressive activity. These immune-active niches cluster tightly in pathway space and reflect the early immune-engaged microenvironment observed in global analyses (Fig. 6). In contrast, epithelial- and fibroblast-rich niches show low EMT, Angiogenesis, and Fibrosis activity at this stage, consistent with minimal structural remodeling in early disease. Together, these patterns indicate that 4-week lesions are dominated by coordinated immune surveillance with limited stromal activation.

**Figure S7.2.**
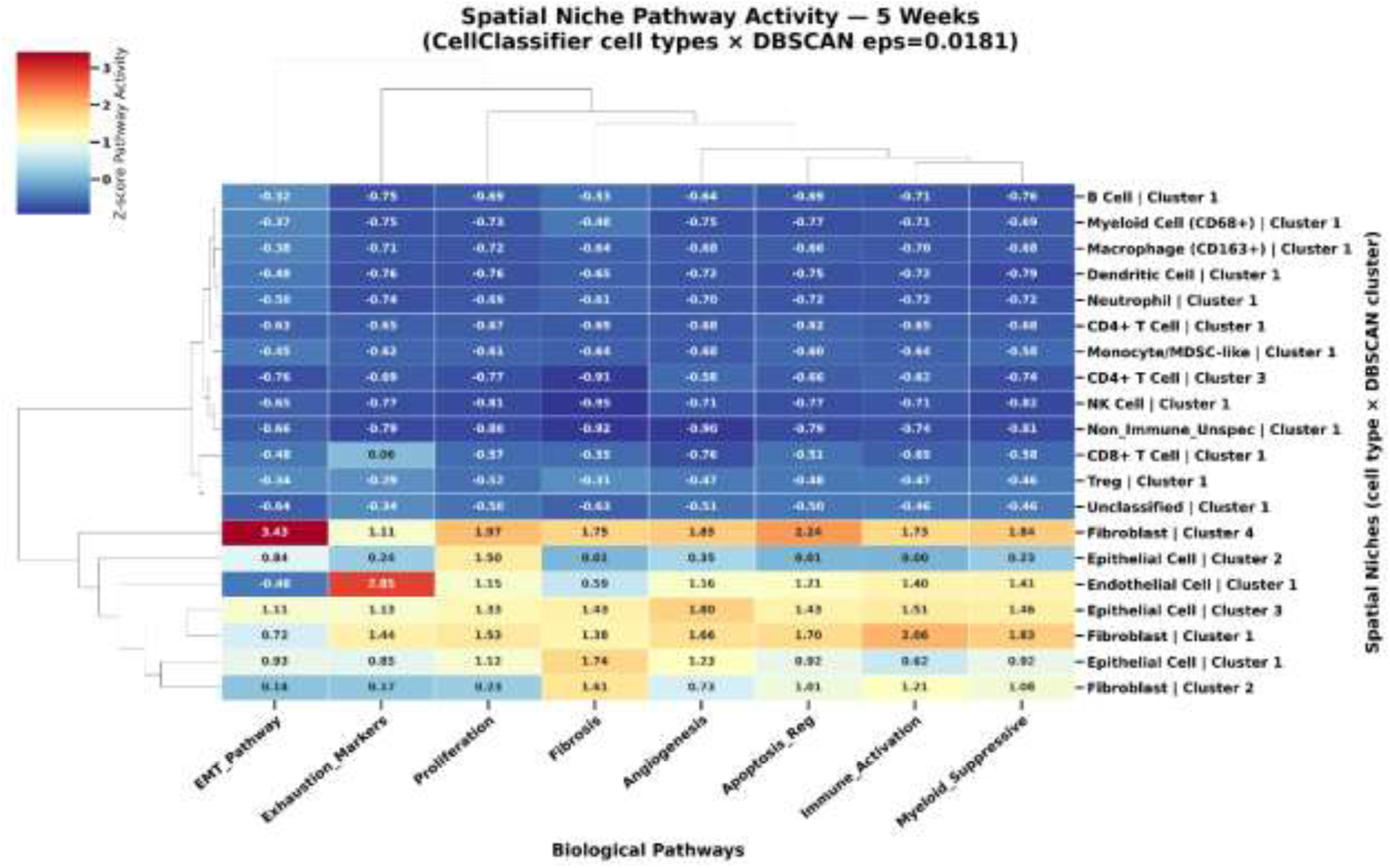
|Spatial niche pathway activity at 5 weeks. Heatmap showing Z-score–normalized pathway activity across spatial niches identified at 5 weeks. Immune-enriched niches—including B cells, CD4⁺ and CD8⁺ T cells, dendritic cells, NK cells, neutrophils, and multiple myeloid subsets—remain prominent and exhibit moderate Immune Activation with low EMT, Angiogenesis, and Fibrosis activity, reflecting a still-immune-engaged microenvironment. Compared with 4 weeks, several immune niches show reduced activation and early signs of pathway diversification, including mild increases in Exhaustion Markers and Myeloid Suppressive activity. Epithelial- and fibroblast-rich niches remain structurally quiescent, with uniformly low EMT and stromal pathway activity. These patterns indicate that 5-week lesions preserve coordinated immune surveillance while beginning a gradual shift toward more heterogeneous immune states.

**Supplementary Figure S7.3.**
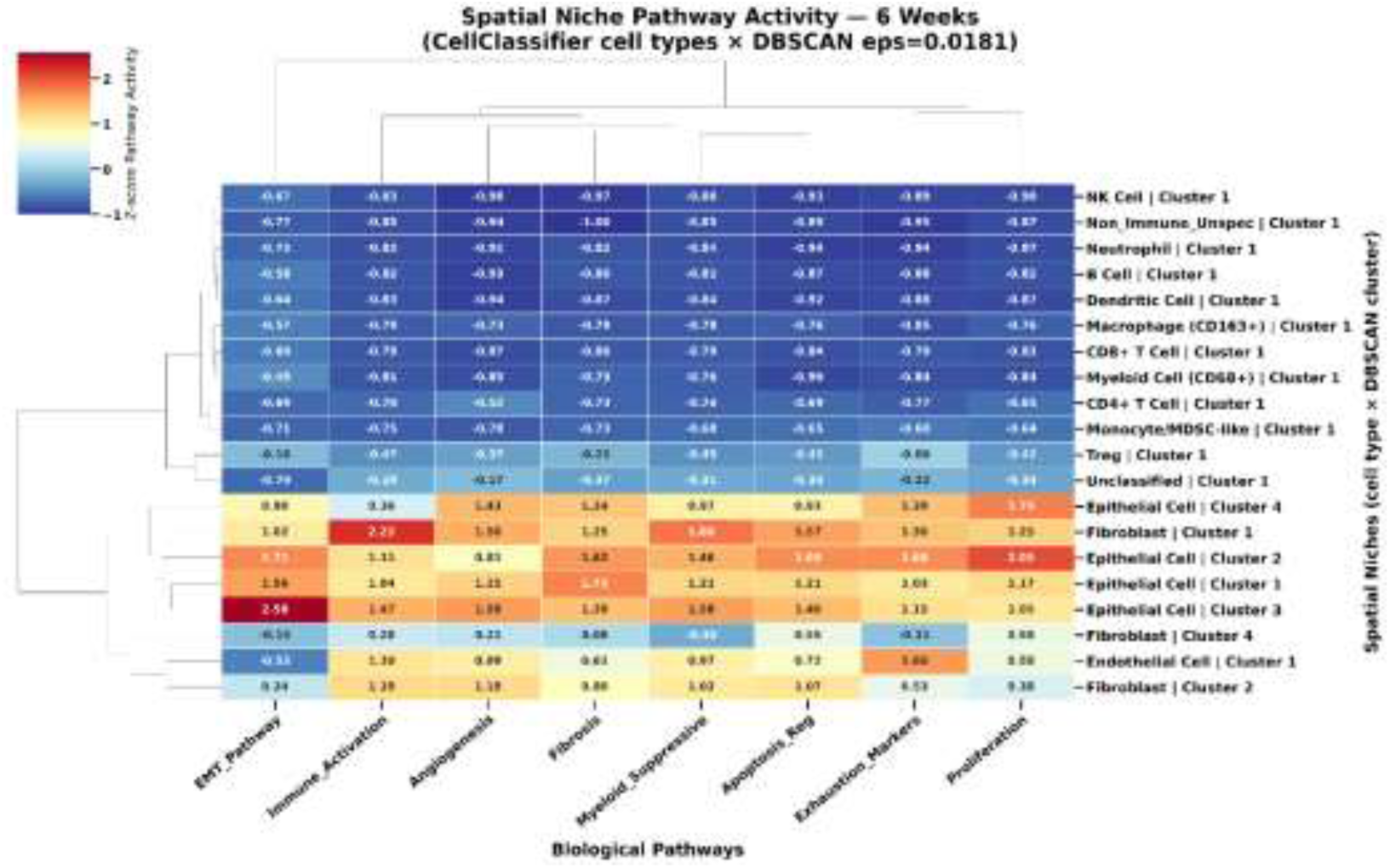
Spatial niche pathway activity at 6 weeks. Heatmap showing Z-score–normalized pathway activity across spatial niches identified at 6 weeks. Immune-enriched niches—including NK cells, dendritic cells, B cells, CD4⁺ and CD8⁺ T cells, neutrophils, and multiple myeloid subsets—remain prominent and continue to exhibit moderate Immune Activation alongside emerging Exhaustion Marker signals. Compared with 4–5 weeks, several immune niches show further attenuation of activation and early divergence in pathway profiles, reflecting the beginning of immune destabilization. Stromal and epithelial niches remain largely quiescent, with uniformly low EMT, Angiogenesis, and Fibrosis activity, indicating that structural remodeling has not yet begun. Overall, the 6-week niche landscape preserves coordinated immune engagement while showing the first signs of transition toward more heterogeneous immune states.

**Supplementary Figure S7.4.**
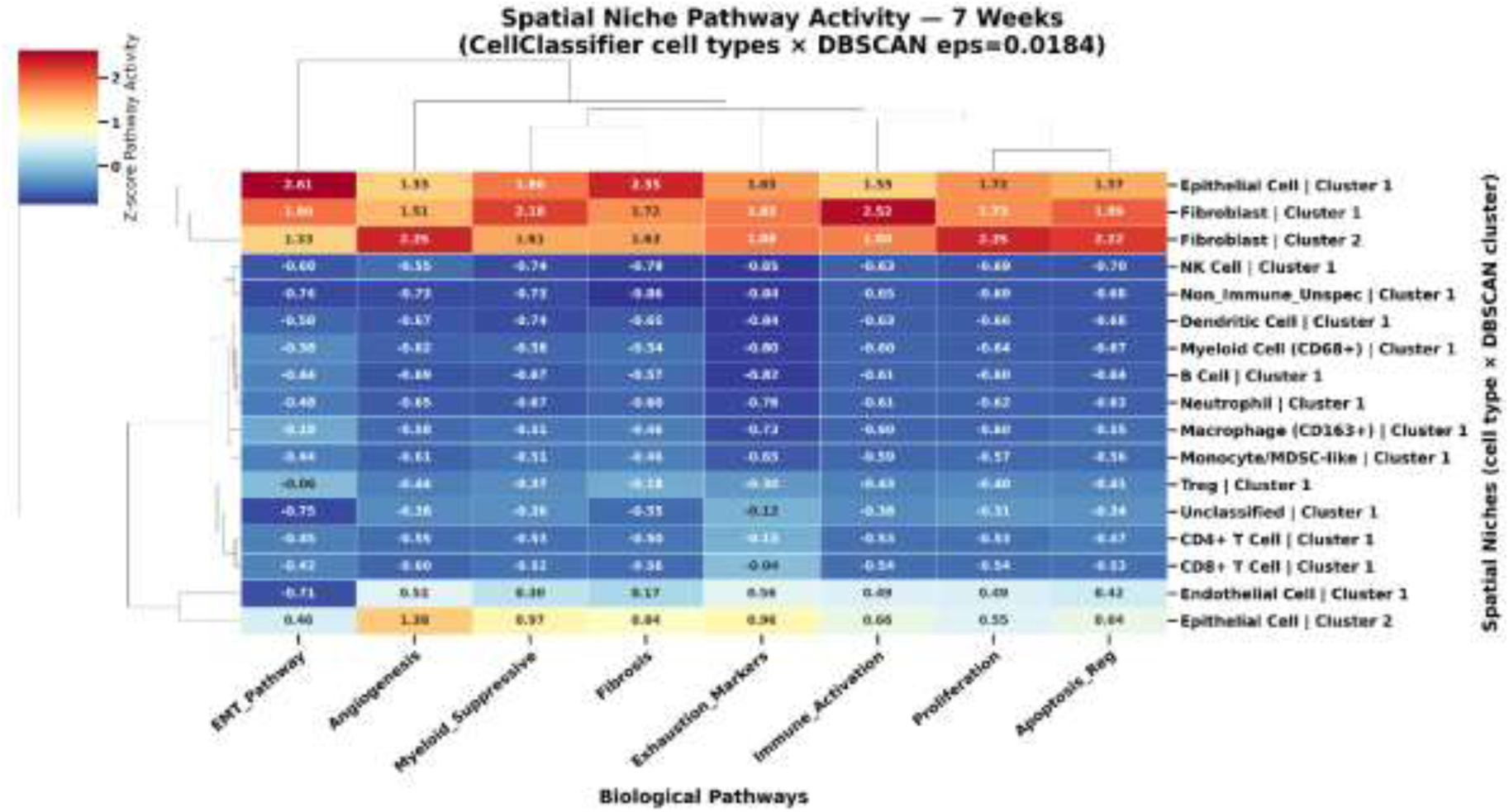
Spatial niche pathway activity at 7 weeks. Heatmap showing Z-score–normalized pathway activity across spatial niches identified at 7 weeks. This stage marks a clear inflection point in microenvironmental organization. Immune-enriched niches—including T cells, B cells, dendritic cells, NK cells, neutrophils, and multiple myeloid subsets—exhibit broad reductions in Immune Activation and increases in Exhaustion Markers, indicating a coordinated decline in lymphoid engagement. In parallel, fibroblast- and epithelial-rich niches begin to display elevated EMT, Angiogenesis, and Fibrosis activity, reflecting the emergence of structurally driven programs. The coexistence of weakened immune niches and increasingly active stromal niches defines a mixed immune–stromal landscape, consistent with the transitional state observed in global pathway dynamics (Fig. 6) and the Petri Net shift toward myeloid-dominated regulation (Fig. 5). These patterns indicate that 7-week lesions occupy a pivotal stage where early immune surveillance gives way to the first signs of stromal remodeling.

**Supplementary Figure S7.5.**
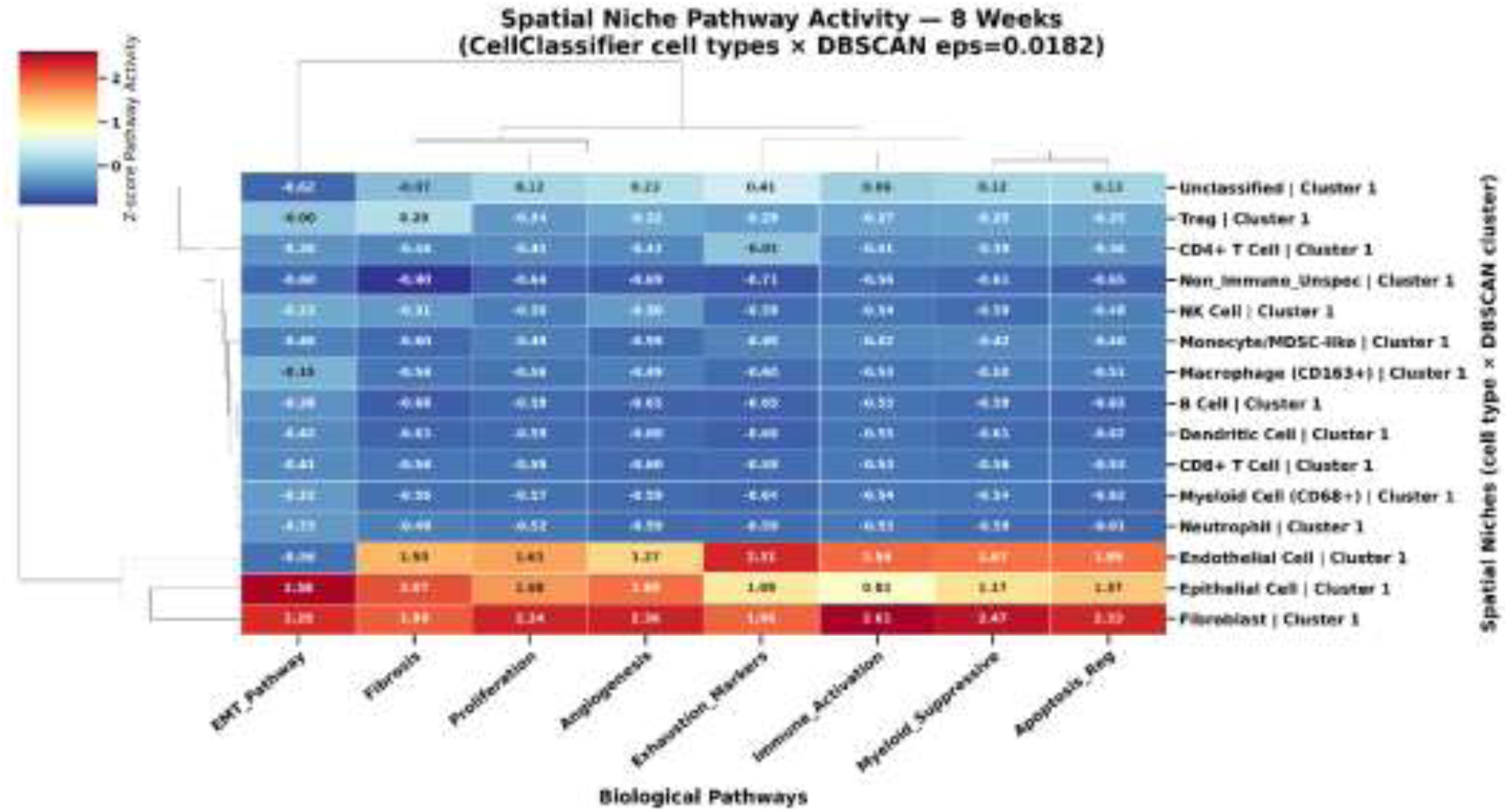
Spatial niche pathway activity at 8 weeks. Heatmap showing Z-score–normalized pathway activity across spatial niches identified at 8 weeks. By this stage, the microenvironment exhibits pronounced divergence between immune and stromal compartments. Immune-enriched niches—including T cells, B cells, dendritic cells, NK cells, neutrophils, and multiple myeloid subsets—show markedly reduced Immune Activation and elevated Exhaustion Markers, indicating substantial attenuation of early immune surveillance. In contrast, fibroblast- and epithelial-rich niches display strong induction of EMT, Angiogenesis, and Fibrosis activity, marking the emergence of structurally driven programs. These stromal-dominant niches cluster tightly in pathway space and reflect the onset of coordinated tissue remodeling. The coexistence of exhausted immune niches and increasingly activated stromal niches defines a transitional microenvironment consistent with the global pathway inflection observed at 8–9 weeks (Fig. 6), signaling the shift toward a tumor-permissive state.

**Supplementary Figure S7.6.**
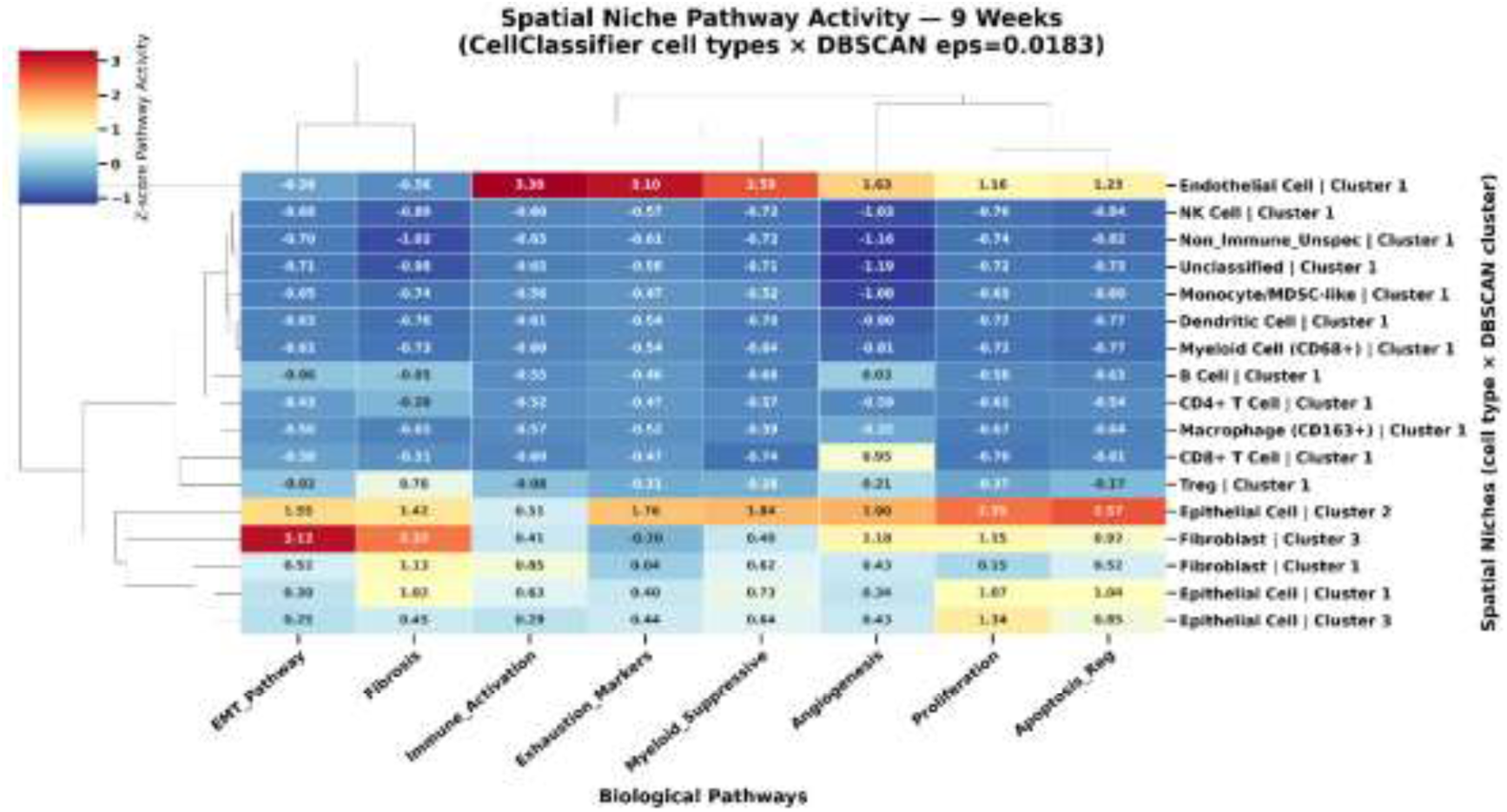
Spatial niche pathway activity at 9 weeks. Heatmap showing Z-score–normalized pathway activity across spatial niches identified at 9 weeks. This stage marks a decisive transition toward stromal-dominated microenvironments. Fibroblast- and epithelial-rich niches exhibit strong induction of EMT, Fibrosis, Angiogenesis, and Proliferation activity, forming a coherent cluster of highly activated stromal programs. In contrast, immune-enriched niches—including T cells, B cells, dendritic cells, NK cells, neutrophils, and myeloid subsets—show minimal Immune Activation and elevated Exhaustion Markers, reflecting profound attenuation of lymphoid engagement. Several immune niches also display increased Myeloid Suppressive activity, consistent with the rise of myeloid-driven regulatory states observed in global analyses (Fig. 6). Together, these patterns indicate that by 9 weeks, the microenvironment has shifted from mixed immune–stromal organization to a predominantly stromal and proliferative state, marking the onset of invasive remodeling.

**Supplementary Figure S7.7.**
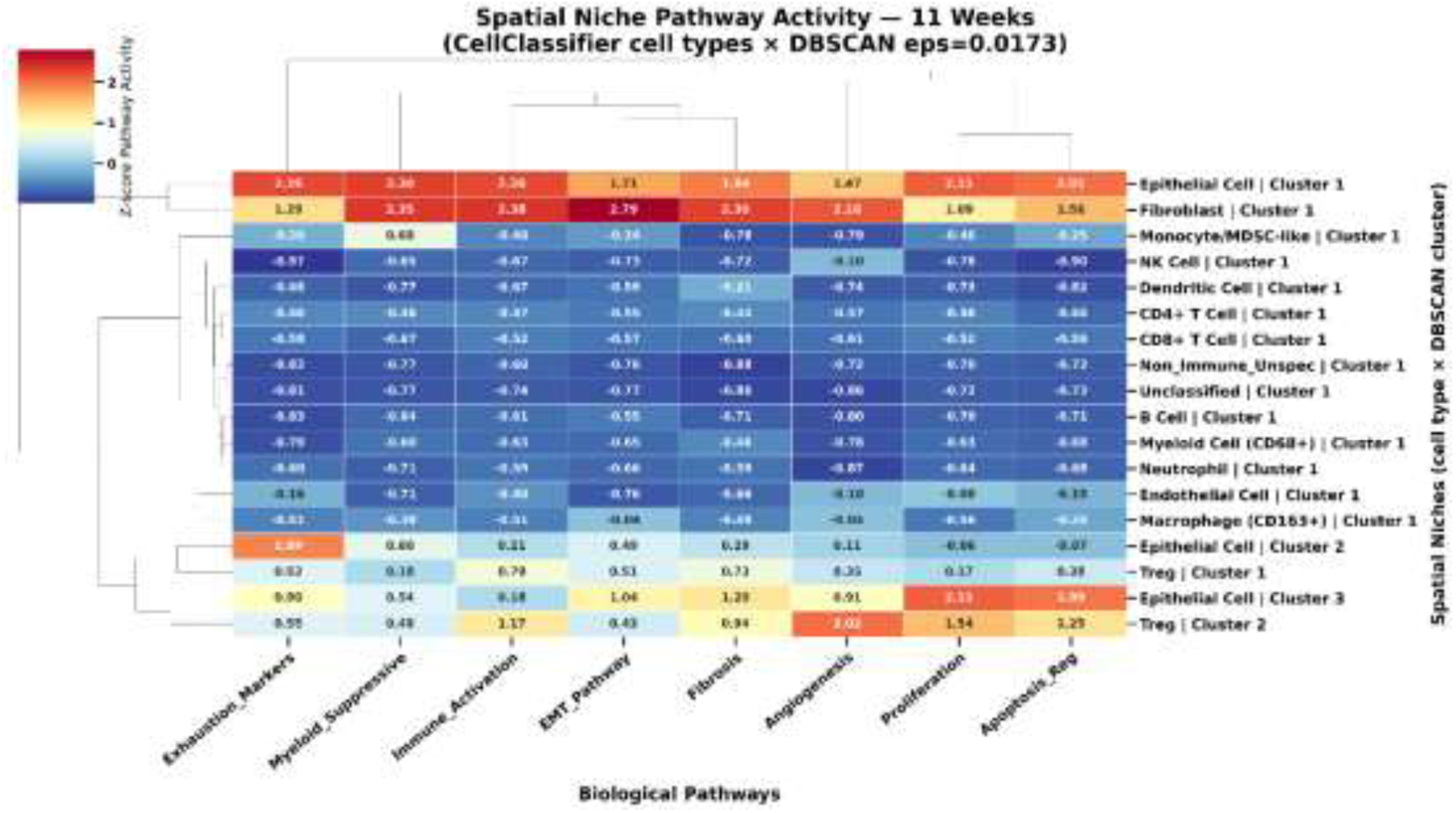
Spatial niche pathway activity at 11 weeks. Heatmap showing Z-score–normalized pathway activity across spatial niches identified at 11 weeks. By this stage, immune-enriched niches—including T cells, B cells, dendritic cells, NK cells, neutrophils, and myeloid subsets—exhibit uniformly low Immune Activation and elevated Exhaustion Markers, reflecting a deeply suppressed immune landscape. Several immune niches also show reduced Myeloid Suppressive activity compared with earlier stages, indicating collapse rather than reorganization of immune function. In contrast, fibroblast- and epithelial-rich niches display moderate activation of EMT, Fibrosis, and Angiogenesis pathways, marking continued structural remodeling despite the downturn in global stromal activity seen at this stage (Fig. 6). The dominance of stromal and epithelial niches, coupled with the near-complete loss of immune activation, indicates that 11-week lesions occupy a late, immunologically silent state preceding the fully invasive stromal expansion observed at 12 weeks.

**Supplementary Figure S7.8.**
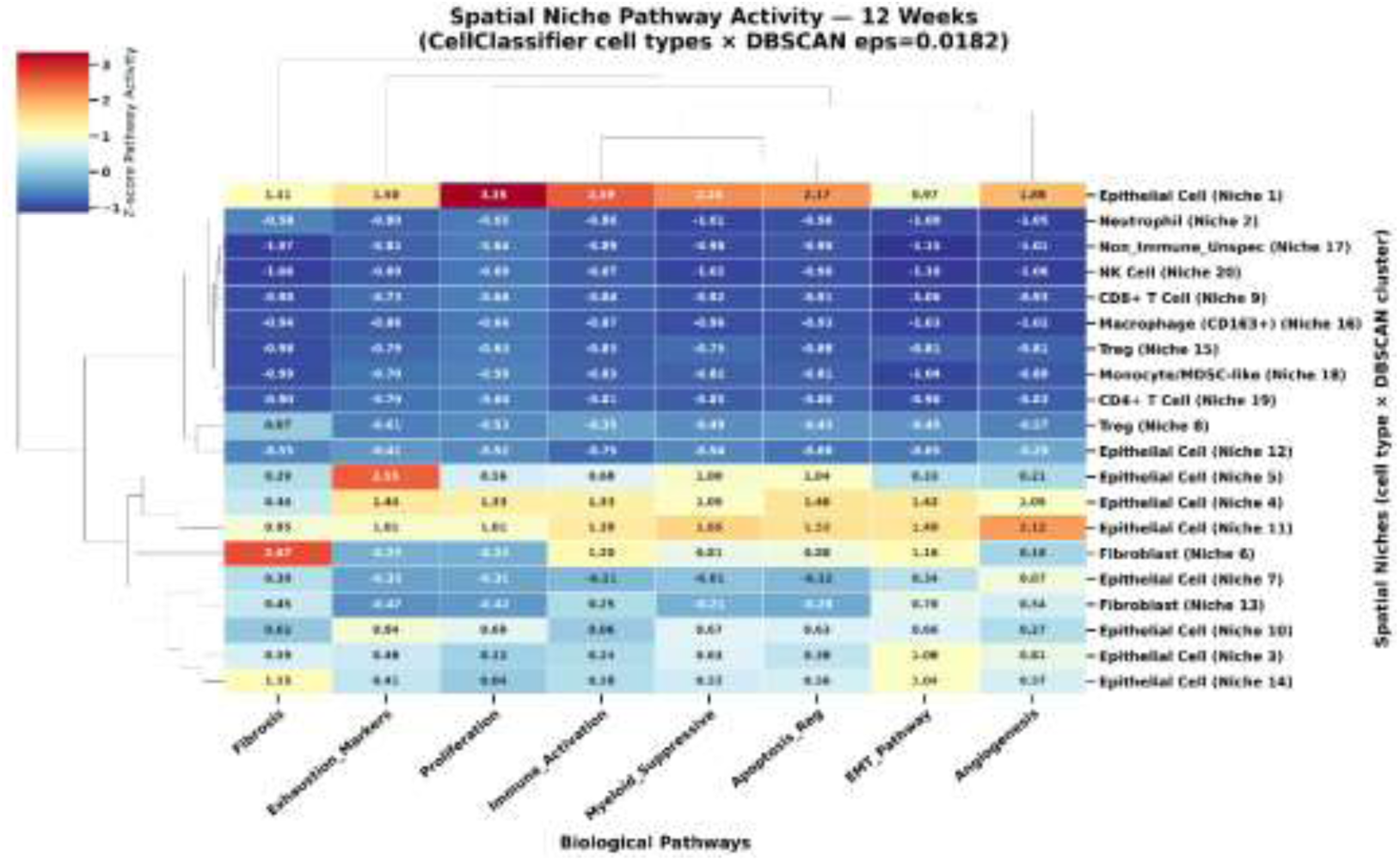
Spatial niche pathway activity at 12 weeks. Heatmap showing Z-score–normalized pathway activity across spatial niches identified at 12 weeks. At this fully advanced stage, the microenvironment is dominated by epithelial- and fibroblast-rich niches exhibiting strong activation of Proliferation, EMT, Angiogenesis, Fibrosis, and Myeloid Suppressive pathways, forming a coherent cluster of highly remodeled stromal and epithelial programs. Immune-enriched niches—including T cells, B cells, dendritic cells, NK cells, neutrophils, and myeloid subsets—show minimal Immune Activation and elevated Exhaustion Markers, reflecting a profoundly suppressed and functionally exhausted immune landscape. Several epithelial niches display exceptionally high Proliferation and Apoptosis Regulation activity, consistent with accelerated epithelial turnover and invasive remodeling. Together, these patterns indicate that by 12 weeks, lesions have transitioned into a fully stromal-dominant, immune-silent state characterized by aggressive epithelial–mesenchymal transition and vascular expansion, marking the culmination of the microenvironmental trajectory observed across earlier stages.

### · Supp. Section 4.6.1 CD68⁺ Myeloid Cell Expansion

Sharma et al. quantified CD68⁺ macrophage infiltration in the tumor stroma by IHC, finding significantly higher stromal CD68⁺ positivity in KSC mice (∼3–4%) compared to controls (<1%; ****P < 0.0001; Fig. 5D) and confirming progressive macrophage recruitment from the earliest postnatal stages. Consistent with this, ROSIE inferred a progressive expansion of CD68⁺ myeloid cells across 4–12 weeks (Fig. 3B), with a log₂ fold change of 1.35 at the 7→8 week transition (Fig. 4)—a period corresponding to the escalation from mPanIN-2/3 to pre-invasive disease. The directional agreement between antibody-based quantification and computational inference supports the fidelity of ROSIE’s myeloid lineage predictions in this model.

### · Supp. Section 4.6.2 Macrophage Activation State and TSPO Expression

V-1520 uptake—a functional readout of TSPO-expressing tumor-associated macrophage (TAM) activity—was quantified at each disease stage: early postnatal (pancreas/muscle ratio 2.3×), adolescence (3.4×), and adulthood (2.7×; Fig. 2B). TSPO immunofluorescence independently confirmed a statistically significant elevation of TSPO protein in adolescence and adulthood relative to controls (**P < 0.01; Supp. Fig. S3). In parallel, ROSIE-inferred Myeloid Antigen Presentation pathway activity peaked at 4 weeks (Z-score = 0.35) and declined progressively through 9–11 weeks (Z = −0.43; Fig. 6), consistent with an early activated myeloid state that transitions toward a suppressive phenotype as disease advances. The concordance between tracer-based functional imaging and ROSIE’s pathway-level inference further supports the accuracy of ROSIE’s myeloid state characterization.

### · Supp. Section 4.6.3 M2 Macrophage Polarization and Immunosuppressive Trajectory

IHC with CD80 (M1 marker) and CD163 (M2 marker) revealed that KSC mice harbor predominantly CD163⁺ M2 macrophages in the tumor stroma, whereas KC mice with pancreatitis exhibit predominantly CD80⁺ M1 macrophages (Supp. Fig. S7). This M2 dominance in KSC tissue indicates a pro-tumorigenic, immunosuppressive myeloid phenotype. ROSIE’s pathway analysis independently captured this shift: T Cell Exhaustion and suppressive myeloid signatures rose progressively from mid-stage onward, with immune-active spatial niches replaced by myeloid-suppressive niches by 11–12 weeks (Fig. 6, Fig. 7). The mechanistic alignment between antibody-defined macrophage polarization and ROSIE’s inferred suppressive programs provides independent biological support for the immune trajectory identified computationally.

### · Supp. Section 4.6.4 Stromal Expansion, Fibrosis, and Acinar Cell Loss

Trichrome-based morphometry (HALO software) in Sharma et al. quantified progressive increases in stromal cell area and collagen deposition, alongside a statistically significant decrease in acinar cell area across 4–12 weeks (****P < 0.0001; Fig. 3C–3D). These tissue-level changes are hallmarks of desmoplastic progression accompanying pancreatic cancer development. ROSIE’s structural inferences mirror these findings: fibroblast-rich cell fractions peak at early-to-mid stages, EMT pathway Z-scores increase to 0.21 at 12 weeks, and Fibrosis activity reaches Z = 1.77 at 12 weeks—among the highest pathway scores detected (Fig. 6). Similarly, ROSIE’s detection of progressive epithelial (ductal) cell expansion is consistent with the histologically confirmed ductal lesion burden quantified by Sharma et al. (***P < 0.001; Fig. 3B). This multidimensional structural concordance validates ROSIE’s ability to capture tissue remodeling dynamics from H&E images alone.

### · Supp. Section 4.6.5 Disease Staging Framework

Histopathologic confirmation of three discrete stages in the same KSC model at identical ages revealed: Early postnatal (4–5 wk, mPanIN-2), Adolescence (6–10 wk, mPanIN-2/3), and Adulthood (11–12 wk, invasive PDAC). Computational analysis applied to the same 4–12 week KSC WSIs demonstrated that ROSIE-identified immune-active ® mixed ® stromal-dominant transitions align precisely with these histopathologic stage boundaries, validating the disease staging framework derived from computational inference.

### · Supp. Table 4.6 Concordance between ROSIE and Wet-Lab Measurements (7 indicators)

**Table 4.6.**
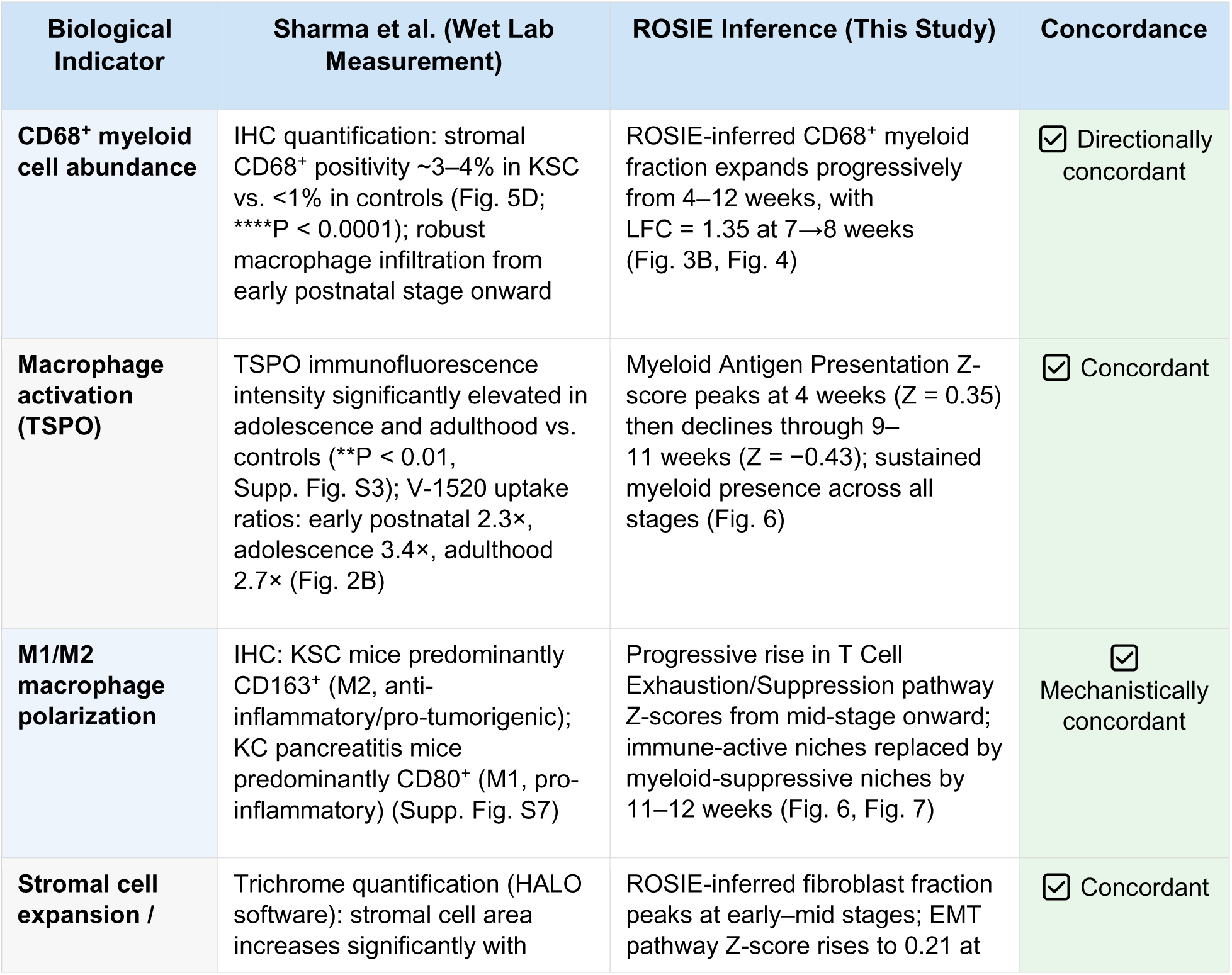
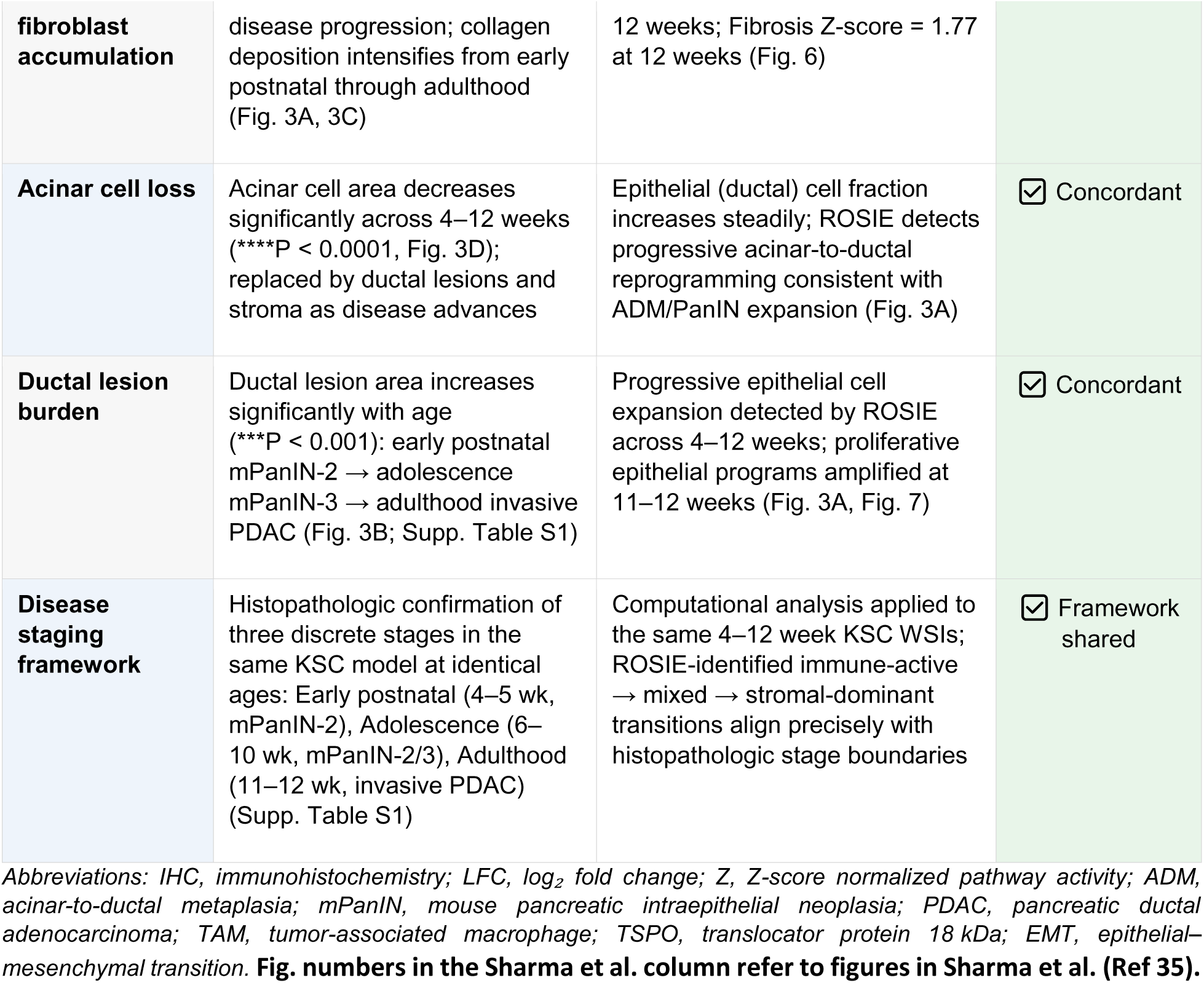
Concordance between ROSIE-inferred predictions and orthogonal wet-lab measurements from Sharma et al. ^35^ **in the KSC mouse model. Supp. Fig. and Supp. Table S1 in the Sharma et al. column also refer to Sharma et al. (Ref 35).**

